# Astrocyte-Specific dsRNA–ADAR1 Signalling Links Innate Immunity to Complement Activation in Parkinson’s Disease

**DOI:** 10.64898/2026.10.06.757209

**Authors:** Yang Liu, Glenda M Halliday, YuHong Fu

## Abstract

**Background:** Dysregulated innate immune activation is a key feature of Parkinson’s disease (PD), but the upstream triggers and cell-type-specific mechanisms remain unclear. Double-stranded RNA (dsRNA) has emerged as a potential driver of innate immune responses. Adenosine deaminase acting on RNA 1 (ADAR1), particularly its interferon-inducible p150 isoform, edits dsRNA to limit aberrant immune activation. Whether this dsRNA–ADAR1 axis contributes to neuroinflammation in PD in a cell-type-specific manner is unknown.

**Methods:** We performed immunofluorescence on post-mortem cingulate cortex tissue from PD cases and controls to localise dsRNA, ADAR1 p150 and pathological α-synuclein protein. We integrated these findings with single-nucleus RNA sequencing data from three independent human brain datasets. An eight-module gene scoring framework was applied to assess dsRNA processing, sensing, and downstream innate immune pathways in neurons and astrocytes.

**Results:** In PD cortex, the proportion of dsRNA-positive astrocytes was higher than in controls, whereas neuronal dsRNA positivity did not differ between groups. ADAR1 p150 positivity increased in both astrocytes and neurons, and the proportion of ADAR1 p150 positive astrocytes correlated with astrocytic α-synuclein burden. Transcriptomic analyses showed that neurons were enriched for dsRNA editing and processing machinery, whereas astrocytes exhibited higher expression of dsRNA sensing pathways, innate immune responses, and transposable element control. While overall pathway activity did not differ between PD and controls after covariate adjustment, network analyses revealed cell-type-specific coupling patterns. Notably, complement gene expression showed a strong and selective association with dsRNA-related pathways in astrocytes, whereas interferon and NF-κB signalling were shared across cell types. In both cell types, ADAR1-associated gene programmes were enriched for oxidative phosphorylation and PD-related pathways.

**Conclusions:** These findings identify astrocytes as a central site of relative dsRNA accumulation and ADAR1 p150 induction in the PD cortex. The selective coupling of dsRNA sensing to complement activation in astrocytes highlights a cell-type-specific innate immune pathway that is constitutively present rather than induced by disease. Together, our protein-level and network-level findings converge on a coherent and testable model in which a constitutive astrocytic circuit is engaged by disease-associated dsRNA accumulation, with potential relevance for disease progression and therapeutic targeting.

## Background

Parkinson’s disease (PD) is the second most common neurodegenerative disorder, characterised by the progressive accumulation of misfolded α-synuclein and selective vulnerability of dopaminergic neurons in the substantia nigra (1, 2). While neuronal loss defines the clinical syndrome, cortical pathology becomes prominent in advanced stages, with α-synuclein inclusions detected in both neurons and astrocytes (3, 4). Astrocytes internalise extracellular α-synuclein through cell-to-cell transfer and endocytosis, and this accumulation drives reactive astrogliosis, mitochondrial dysfunction, and innate immune activation (5–7).

Neuroinflammation is increasingly recognised as a central feature of PD pathogenesis rather than a secondary consequence of neuronal death (8, 9). In the PD brain, pathological α-synuclein activates astrocytes and microglia through NF-κB, NLRP3 inflammasome, and toll-like receptor (TLR) signalling pathways, driving release of IL-1β, IL-6, and TNF-α (8, 10). Emerging evidence positions astrocytes as active contributors to this inflammatory cascade. Astrocytic uptake of α-synuclein engages both TLR4-dependent and independent mechanisms, promoting cytokine release and non-cell-autonomous neuronal injury (11, 12). In a viral encephalitis model of parkinsonism, astrocyte-specific NF-κB signalling activation is sufficient to drive dopaminergic neuronal loss and α-synuclein aggregation (13). In human induced pluripotent stem cell (iPSC)-derived astrocytes from PD patients carrying *LRRK2* mutations, transcriptomic profiling reveals upregulation of antigen presentation, complement activation, and IL-6 signalling pathways, while astrocyte-derived IL-6 drives STAT3-mediated neuronal death that can be rescued by IL-6 receptor blockade (14). However, the upstream molecular triggers that drive these changes in PD brain remain incompletely understood.

One upstream candidate that may trigger such a response is endogenous double-stranded RNA (dsRNA), an emerging danger-associated molecular pattern (DAMP) molecule recognised by the cytosolic sensors MDA5 (encoded by IFIH1) and RIG-I (encoded by DDX58) (15, 16). Endogenous dsRNA arises primarily from bidirectional transcription of repetitive elements, including Alu elements and endogenous retroviruses (ERVs), and must be edited by adenosine deaminases acting on RNA (ADAR) to prevent aberrant immune activation (17, 18). Garza *et al.* recently demonstrated region- and cell-type-specific transposable element (TE) activation linked to interferon responses in PD, with the strongest signal in substantia nigra microglia (19), leaving open whether dsRNA pathway dysregulation extends to additional regions and cell types through mechanisms beyond bulk TE expression.

ADAR1, particularly its IFN-inducible p150 isoform, is the principal editor that destabilises endogenous dsRNA and prevents MDA5-dependent type I IFN signalling (18, 20–23). Loss of function mutations in ADAR1 leads to Aicardi-Goutières syndrome, a severe interferonopathy that underscores the critical role of ADAR1-mediated editing in immune homeostasis (24). Increasing evidence implicates the dsRNA–ADAR1 axis in neurodegeneration. D’Sa *et al.* demonstrated that α-synuclein oligomers trigger ADAR1 upregulation and isoform switching in human iPSC-derived astrocytes, accompanied by increased adenosine-to-inosine (A-to-I) RNA editing enrichment in inflammatory pathways, and elevated editing in astrocytes in postmortem PD brain (25). Similarly, ADAR1 knockdown in primary human astrocytes triggers type I IFN pathway activation and accumulation of TE transcripts with dsRNA-forming potential (26). Altered ADAR1 expression and editing patterns that have also been reported in Alzheimer’s disease and amyotrophic lateral sclerosis (27, 28). Together, these findings establish the dsRNA–ADAR1 axis as a plausible upstream regulator of innate immune activation in neurodegeneration.

Despite this growing evidence, prior work has examined RNA editing activity (25) or TE expression (19) largely in isolation and not addressed whether the coupling between dsRNA processing, sensing, and downstream immune effector programmes is organised in a cell-type-specific manner in the PD brain. Whether astrocytes and neurons differ not only in absolute expression of dsRNA pathway components but in how tightly these components are co-regulated with downstream effectors has not been investigated. In this study, we combined immunofluorescence analysis of post-mortem PD and control cingulate cortex tissue with single-nucleus RNA-sequencing (snRNA-seq) from three publicly available datasets. Specifically, we (1) compare the proportion of dsRNA- and ADAR1 p150-positive cells between neurons and astrocytes in PD and control cingulate cortex; (2) define the cell-type-specific transcriptomic architecture of the dsRNA–ADAR1–innate immune axis using an eight-module gene scoring framework; (3) determine whether dsRNA sensing differentially couples to innate immune effector programmes in astrocytes and neurons; and (4) identify functional pathways associated with dsRNA pathway activation in astrocytes. We propose a neuron-astrocyte division-of-labour model in which neurons sustain a high throughput dsRNA processing and editing, while astrocytes function as the principal sensing and effector compartment for dsRNA-driven innate immune activation in the PD cortex.

## Methods

### Study design and overview

This study integrated immunofluorescence in postmortem human brain tissue with computational re-analysis of publicly available snRNA-seq datasets to interrogate dsRNA biology, ADAR1-mediated A-to-I editing and innate immune signalling in PD. The immunofluorescence arm comprised 44 cases of the cingulate cortex, including sporadic PD (n=23), age-matched controls (n=18), and *SNCA* mutation carriers (n=3), and was used to determining cell types containing dsRNA, ADAR1 p150, and α-synuclein at a single-cell level. The transcriptomic arm comprised 43 donors of the cingulate and frontal cortex (PD, n = 22; Control, n = 21) compiled from three GEO datasets (GSE202210, GSE178146, and GSE253462). An eight-module gene-scoring framework was applied to dissect cell-type-specific pathways in astrocytes and neurons (Fig. 1).

**Figure 1.**
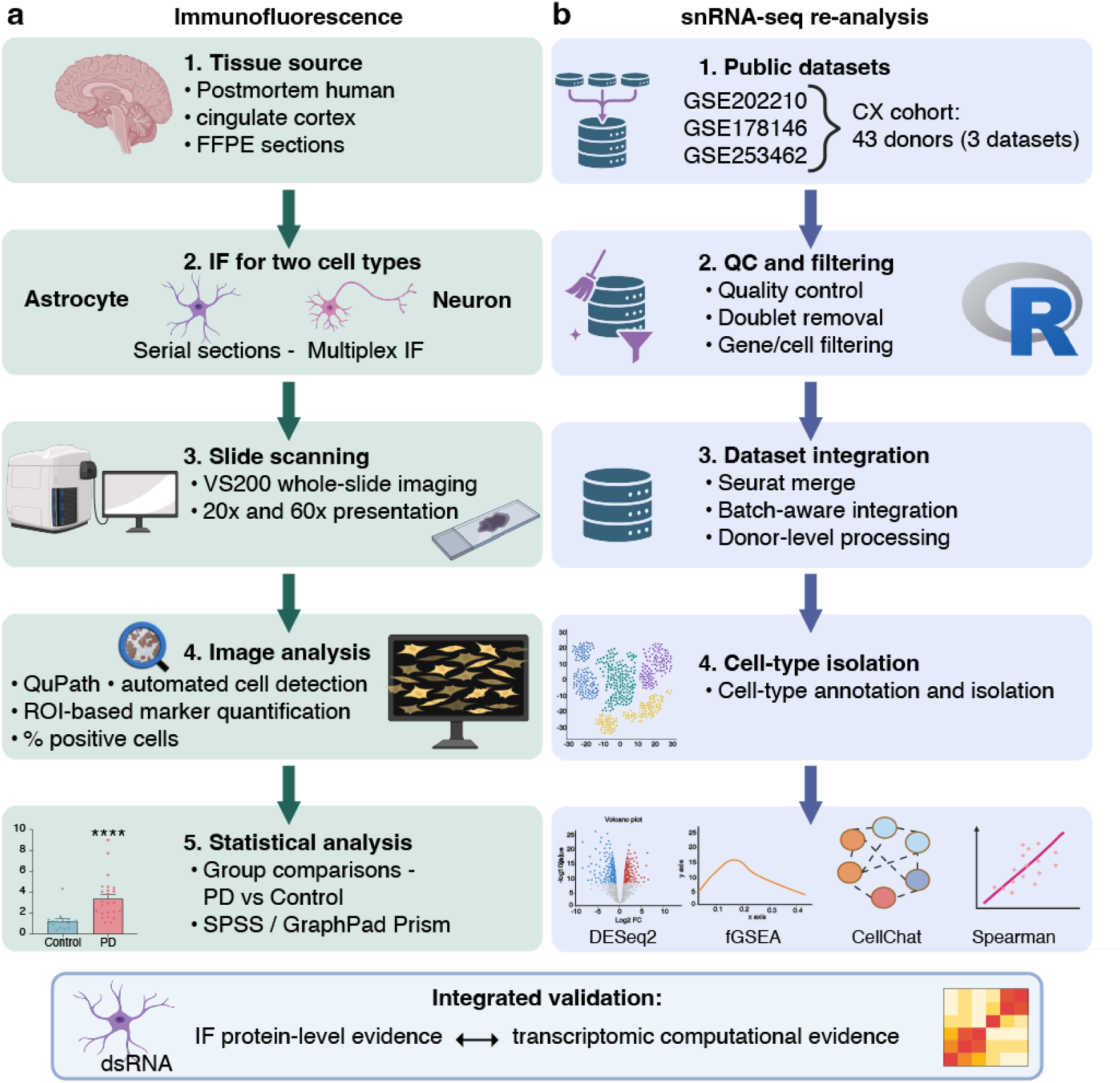
Study design overview. (a) immunofluorescence (IF) strategy for postmortem human cingulate cortex (cohorts in Table 1). (b) snRNA-seq analysis pipeline. Three cortical snRNA-seq datasets (Table 1) were processed under a unified quality-control and integration pipeline to isolate astrocyte and neuron populations. An eight-module gene-scoring framework was applied to quantify dsRNA processing, sensing, transport, innate immune activity, ADAR1 and ADAR2 expression, and transposable element control at single-cell resolution. Abbreviations: CX=cortex, FFPE=formalin-fixed, paraffin-embedded, QC=quality control, ROI-region of interest, PD=Parkinson’s disease. Figure created in BioRender (BioRender.com).

**Table 1.** Case demographics.

| Study - Source | group | Sex (M/F) | Age at death (years) | Postmortem delay (hours) | Disease duration (years) | Braak Lewy stage/6<sup>(3)</sup> (n/stage) |
| --- | --- | --- | --- | --- | --- | --- |
| IF- SBB | Control | 9/9 | 86±8.5 | 21±10 | - | 0 |
| IF- SBB | PD | 16/7 | 77.5±6 ** | 12±9* | 11±2.5 | 13/stage 4, 10/stage 6 |
| IF- SBB | *SNCA* | 3/0 | 56±19** | 32±14 | 13±5 | 3/stage 4-5 |
| RNA - GSE202210<sup>(31, 32)</sup> | Control | 3/3 | 70.0±6.3 | 10.1±4.9 | - | NA |
| RNA - GSE202210<sup>(31, 32)</sup> | PD | 3/3 | 81.8±8.3* | 7.6±2.8 | 17.4±8.4 (n=5) | NA |
| RNA - GSE178146<sup>(33)</sup> | Control | 6/1 | 75.1±13.5 | 18.0±9.3 | - | 0 |
| RNA - GSE178146<sup>(33)</sup> | PD | 2/5 | 83.3±6.3 | 17.7±5.8 | 14.1±4.1 | 2/stage 4-5, 5/stage 6 |
| RNA - GSE253462<sup>(34)</sup> | Control | 5/3 | <65-85 | NA |  | 0 |
| RNA - GSE253462<sup>(34)</sup> | PD | 6/3 | <65-80 | NA | NA | 3/stage 4-5, 6/stage 6 |

Values are mean ± SD. Asterisks denote significant differences compared to controls (\* $p < 0.05$ , \*\* $p < 0.001$ ; one-way ANOVA with post-hoc protected t-tests). Abbreviations: IF=immunofluorescence, F=female, M=male, n=number, PD=Parkinson's disease, SBB=Sydney Brain Bank, *SNCA*= *SNCA* gene mutation.

### Human brain tissue samples

For our immunofluorescence study, formalin-fixed paraffin-embedded cingulate cortex sections (6–8 μm) along with demographic and neuropathological staging were obtained from the Sydney Brain Bank (www.sbb.neura.edu.au), which collects and characterizes cases from regional brain donor programs. Informed consent for banking was acquired by the Sydney Brain Bank under the University of NSW HREC0937. All cases with PD were levodopa-responsive and fulfilled the UK Brain Bank Clinical Criteria for a diagnosis of clinical PD (29) with no other neuropathologies. Three cases were identified with *SNCA* mutations as previously reported (30). The assessment of postmortem human brain tissue was approved by the Human Research Ethics Committee at the University of Sydney (2020/HE000707). Demographic and clinical details are provided in Table 1.

For our transcriptomic analyses, snRNA-seq data from post-mortem human cingulate (or frontal) cortex were obtained from three publicly available datasets in the Gene Expression Omnibus (GEO): GSE202210, GSE178146, and GSE253462. Demographic covariates (age at death, sex, post-mortem delay) and α-synuclein Braak Lewy staging (3) were extracted from the source publications and harmonised across studies (Table 1).

### Immunofluorescence staining and imaging

Multiplex immunofluorescence was performed on 6–8 μm formalin-fixed paraffin-embedded cortical sections to co-localise dsRNA pathway components with cell-type markers and α-synuclein pathology. Sections were deparaffinised, through graded xylene and ethanol, rehydrated and subjected to heat-induced epitope retrieval in 0.01 M sodium citrate buffer (pH 6) for 2 hours. Non-specific binding was blocked with Fc Receptor Blocker (INNOVEX Bioscience, NB309) for 30 minutes at room temperature. Sections were incubated with primary antibodies for 48 hours at 4°C (Table 2).

**Table 2.**
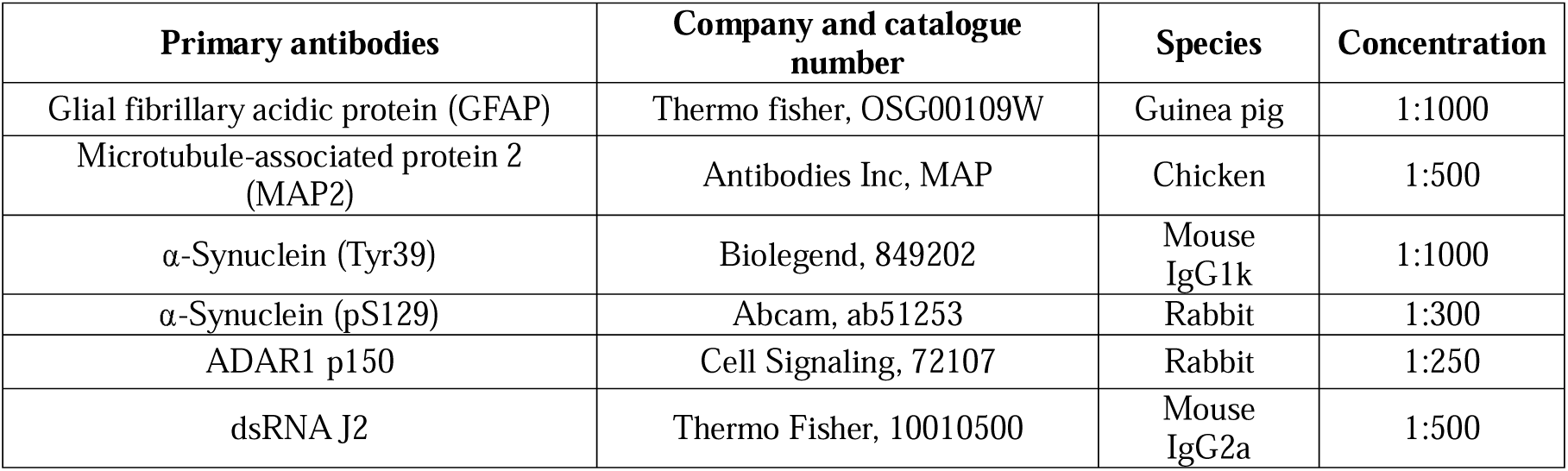
Primary antibodies for immunofluorescence.

Sections were washed and incubated with the secondary antibodies for 2 hours at room temperature (Supplementary Information Table 1). For panels in which ADAR1 p150 was co-detected with α-synuclein pathology markers raised in the same host species, ADAR1 p150 antibody was directly labelled using the Zenon Rabbit IgG Labelling Kit (Thermo Fisher, Z25302) per the manufacturer’s instructions to circumvent host-species cross-reactivity. Nuclei were counterstained with Hoechst 33342 (1:1000). Negative controls consisted of (i) primary antibody omission with secondary only, and (ii) isotype-matched IgG substituted at the same concentration as the primary, both were processed in parallel for each batch and used for intensity-threshold calibration.

Whole-slide fluorescence images were acquired on an Olympus VS200 slide scanner at 20× magnification for quantification and 60× for representative images. Image analysis was performed in QuPath v0.6.0. Cell detection was performed on the DAPI channel. Per-channel marker positivity was determined by intensity thresholding calibrated against the negative-control and secondary-only sections. For each section, a region of interest restricted to cortical grey matter was annotated manually, using neuronal and astrocytic morphology together with the DAPI nuclear pattern to define the grey–white matter boundary. The percentage of marker-positive cells per region of interest was calculated for each case and exported for downstream statistical analysis.

### Statistical analysis of immunofluorescence data

For each immunofluorescence marker (or marker combination), the percentage of positive cells was calculated for each donor individually, yielding one value per donor per marker for downstream statistical analysis. All image analysis and quantification were performed blinded to diagnostic group. Group comparisons were performed by univariate general linear modelling with age, sex and postmortem delay as covariates. *SNCA* mutation carriers were excluded from all quantitative comparisons. Associations between marker-positive proportions were assessed by Spearman rank correlation on covariate-adjusted residuals across the 41 sporadic PD and control cases. Statistical analyses were performed in IBM SPSS Statistics (IBM Corp., Armonk, NY, USA) and GraphPad Prism v11.0 (GraphPad Software Inc.) was used for visualisation.

### Single-nucleus RNA sequencing data processing

Raw count matrices from GSE202210, GSE178146, GSE253462 were processed through a standardised quality control (QC), normalisation, and integration pipeline in R (v4.5.2) using Seurat v5 (35) to minimise systematic inter-dataset bias. Identical filtering thresholds were applied across the three studies (Supplementary Information Table 2). Batch effects were corrected using Harmony (36) applied to the first 30 principal components, with the dataset identifier as the grouping variable. Uniform manifold approximation and projection (UMAP) embedding and shared nearest-neighbour clustering (resolution = 0.5) were computed on the Harmony-corrected dimensions (Supplementary Information Fig. 1).

Cell-type annotation used Seurat’s AddModuleScore (35) with canonical marker gene panels: astrocytes (*AQP4, GFAP, SLC1A2, ALDH1L1, S100B, GJA1*), excitatory neurons (*SLC17A7, CAMK2A, NRGN, SATB2, CUX2*), and inhibitory neurons (*GAD1, GAD2, SLC32A1, SST, PVALB, VIP*). Each nucleus was assigned to the cell type with the maximal marker-module score (Seurat AddModuleScore, control set n = 50). Downstream astrocyte and neuronal populations were defined by two confidence thresholds (highest score > 0.1 and ≥1.5-fold greater than the second highest score). Nuclei failing either criterion were excluded from cell-type-specific analyses. The primary analyses pool excitatory and inhibitory neurons into a single “total neuron” population to maximise statistical power for cell-type contrasts, with separate sensitivity analyses for excitatory and inhibitory neurons reported in Supplementary Datasheets. Astrocytes and total neurons were extracted as separate objects and independently re-clustered with Harmony batch correction and saved as independent Seurat objects prior to all downstream analyses.

### Donor-level covariate adjustment and multiple testing correction

Unless otherwise stated, all donor-level analyses were adjusted for age at death, sex, and postmortem delay as covariates (Table 1) using ordinary least squares regression. Postmortem delay metadata were not available for GSE253462 donors; for these donors postmortem delay was imputed as the cross-cohort mean prior to model fitting. Analyses relying on within-donor pairing (astrocyte versus neuron contrasts) are not covariate-adjusted. Multiple-testing correction used the Benjamini–Hochberg (BH) procedure was applied within each analytical family, with the family scope specified per analysis. Significance was set at BH-adjusted *P* < 0.05.

### Eight-module gene scoring framework

An eight-module framework was designed to interrogate distinct functional nodes of the dsRNA–ADAR1–innate immunity axis. The modules comprised three single-gene anchors and five multi-gene pathway modules: (1) ADAR1 (*ADAR*), (2) ADAR2 (*ADARB1)*, (3) SNCA (*SNCA*), (4) dsRNA processing (*ILF2, ILF3, STRBP, ZFR, DHX9, DDX39B, SFPQ, NONO*; 8 genes (37, 38)), (5) dsRNA/EV (extracellular vesicle) transport (*SIDT1, SIDT2, LAMP2, RAB27A, RAB27B, RAB35, SMPD3, CD63, CD9, CD81, PDCD6IP, TSG101, SDCBP, MVP;* 14 genes (38–42)), (6) dsRNA sensing (*DDX58, IFIH1, DHX58, TLR3, MAVS, TICAM1, TBK1, IKBKE*; 8 genes (16, 43, 44)), (7) innate immune response (*JAK1, TYK2, STAT1, STAT2, IRF9, RELA, ISG15, USP18, OAS1, OAS2, OAS3, OASL, RNASEL, EIF2AK2, PRKRA, MX1, IFIT1, IFIT2, IFIT3, C3,C4A* ; 21 genes (45–47)), and (8) TE control (*MOV10, TRIM28, SETDB1, DNMT1, DICER1, AGO2, L1TD1, PIWIL1*; 8 genes (48–50)).

Per-cell module scores for these eight functional modules were computed as the mean log-normalised expression of constituent genes present in the cell’s expression matrix using Seurat v5 GetAssayData, and for single-gene anchors the score reduces to the log-normalised expression of the respective gene. Cell level scores were then aggregated to donor-level means within each cell type.

### Differential expression and module-level analyses

Gene-level PD and control differential expression analysis was performed using a pseudobulk approach: raw UMI counts were summed per donor per cell type and analysed using DESeq2 (v1.46) with default size-factor normalisation (genes detected in fewer than 10 nuclei were filtered prior to model fitting). *P*-values were adjusted by the BH method. Module-level comparisons between PD and control donors were tested separately in astrocytes and total neurons, using ordinary least-squares models containing diagnosis and the prespecified covariates, with BH correction across the eight modules within each cell type. Unadjusted Wilcoxon rank-sum tests are reported as supplementary comparisons. Within-donor cell-type contrasts (astrocyte versus neuron) were assessed using two-sided paired Wilcoxon signed-rank tests across all 43 donors, with each donor contributing matched astrocyte and neuron module scores, with BH correction across the eight modules.

### Cross-module coupling analysis

Donor-level relationships between modules were quantified using partial Spearman correlations within each cell type (ppcor v1.1), adjusting for age at death, sex and postmortem delay. P-values were BH-corrected across all cell type, disease group and module-pairs.

### Innate immune sub-panel analysis

For all coupling analyses, the composite innate immune module was replaced by nine mechanistically interpretable sub-panels, while the remaining seven modules were retained unchanged; module-sub-panel analyses therefore involve seven modules and nine sub-panels. Sub-panels were curated and scored independently in astrocytes and neurons (Supplementary Information Table 3): ISG core (12 genes); IFN–STAT (8 genes); NF-κB (10 genes); complement (9 genes); OAS–RNase L (5 genes); astrocyte reactivity (6 genes); inflammasome (9 genes); TLR–dsRNA (3 genes); and TLR–DAMP (7 genes).

Donor-level sub-panel scores were computed using Seurat’s AddModuleScore approach with 25 expression bins and 100 control genes per panel gene, then averaged across cells per donor. Module and sub-panel coupling were assessed within each cell type as partial Spearman correlations. Astrocyte and neuron differences were evaluated using 10,000 paired donor-level bootstrap resamples; for this comparison we report the difference in correlation on the native scale, Δρ = ρ_astrocyte − ρ_neuron, with percentile 95% confidence intervals. P-values were by referring z = Δρ / SE_bootstrap to the standard normal distribution and were obtained from the standard normal distribution and BH-adjusted across all module pairs.

### Functional gene set enrichment analysis

For each cell type and module, we constructed cell-level ranked gene lists by computing the Spearman rank correlation of each gene’s log-normalised expression with the donor-covariate-residualised module score across all cells in that cell type. Functional gene set enrichment analysis (fGSEA) was performed with the fgsea package (v1.32.4) using nPermSimple = 10000, minSize = 15, and maxSize = 500, against KEGG human pathway gene sets retrieved from MSigDB (v2024.1.Hs) via msigdbr. *P*-values were BH-adjusted within each cell type and module analysis.

### Reactive astrocyte gene-signature analysis

Reactive-astrocyte transcriptional signatures were assessed using three canonical marker panels derived from Liddelow *et al.* (51): the A1-associated panel (*C3, SERPING1, GBP2, FKBP5, PSMB8, SRGN, AMIGO2, C1R, C1S, CFB, CFI;* 11 human-mappable genes after exclusion of murine-specific H2-T23 and GGTA1), the A2-associated panel (*EMP1, S100A10, SLC10A6, TM4SF1, CD109, PTX3, SPHK1, CD14, TGFB2, B3GNT5*; 10 genes), and a pan-reactive panel (*GFAP, VIM, SERPINA3, LCN2*; 4 genes). The A1-associated panel was additionally divided into complement and non-complement subsets, generating A1-all, A1-complement (*C3, C1R, C1S, CFB, CFI, SERPING1)*, and A1-non-complement (*FKBP5, GBP2, PSMB8*) composite scores. SRGN and AMIGO2 were retained only in the A1-all composite because they belong to neither functional subset. This subdivision was used to distinguish complement-enriched from non-complement components of the historically A1-associated signature. Composite scores were calculated from donor-level astrocyte pseudobulk profiles using the same scoring procedure as the functional gene modules. Each of the five reactive-astrocyte signature scores was then tested for association with the seven prespecified functional module scores using Spearman correlations. Diagnostic-group-stratified correlations and the descriptive group difference Δρ = ρ_PD − ρ_Control are reported for the A1-complement composite. Multiple-testing correction was performed across the 35 signature–module comparisons using the BH test.

### Cell-type specific divergence across α-synuclein Braak stages

To assess whether the cell-type divergence in dsRNA model activity changes with α-synuclein pathology severity, we used the subset of 31 cortical donors with available α-synuclein Braak staging from GSE178146 and GSE253462: 15 controls at stage 0, 5 PD cases at stages 4–5 (Braak 4, n =1, Braak 5, n= 4), 11 PD cases at stage 6. GSE202210 was excluded because Braak metadata were not available. Donor-level module scores were computed as described above for astrocytes, excitatory neurons, inhibitory neurons, and total neurons. Within-donor divergence was defined as the astrocyte module score minus the matched neuron module score, which cancels donor-specific technical variation that affects both cell types equally.

The associations between within-donor divergence and Braak stage groups was estimated by ordinary least squares with age and dataset of origin as covariates. Braak stage groups were encoded as an ordinal numeric variable (0 = control, 1 = Braak 4–5, 2 = Braak 6). Point estimates and 95% confidence intervals were obtained from 10,000 donor-level bootstrap resamples, and P values were BH-corrected across the eight modules within each cell type. Given that only five donors fell in the Braak 4–5 group, this analysis is underpowered and its null results are reported as uninformative rather than as evidence of stability in Supplementary Datasheets.

### Software and code availability

All analyses were performed in R v4.5.2 with Bioconductor and CRAN packages. Key packages included Seurat v5 and SeuratObject for snRNA-seq handling, Harmony for batch correction, scDblFinder for doublet detection, DESeq2 v1.46 for pseudobulk differential expression, ppcor for partial correlations, fgsea v1.32.4 with msigdbr for gene-set enrichment analysis. All analysis code is available at https://doi.org/10.5281/zenodo.21315999.

## Results

### dsRNA accumulates selectively in astrocytes correlating with astrocytic α-synuclein accumulation

To characterise dsRNA presence in the PD brain, we performed immunofluorescence using the J2 antibody (green puncta, Fig. 2a,c), which specifically recognises dsRNA duplexes of 40 base pairs or longer (52). In cingulate cortex sections, the proportion of GFAP-positive astrocytes with J2 dsRNA puncta (Fig. 2a,b) was significantly higher in PD than in controls (11.27 ± 1.15% vs 6.29 ± 1.41%, *P* = 0.019, Fig. 2e), whereas the proportion of J2-positive neurons (Fig. 2c,d) did not differ between groups (4.87 ± 0.59% vs 3.89 ± 0.76%, *P* = 0.273; Fig. 2f). However, the proportion of cells positive for ADAR1 p150 was significantly elevated in both cell types in PD cases (magenta puncta, Fig. 2a-d, astrocytes (9.55 ± 0.70% vs 5.93 ± 0.85%, *P* = 0.006, Fig. 2e and neurons 6.91 ± 0.66% vs 2.74 ± 0.85%, *P* = 0.002, Fig. 2f), indicating activation across cell types.

**Figure 2.**
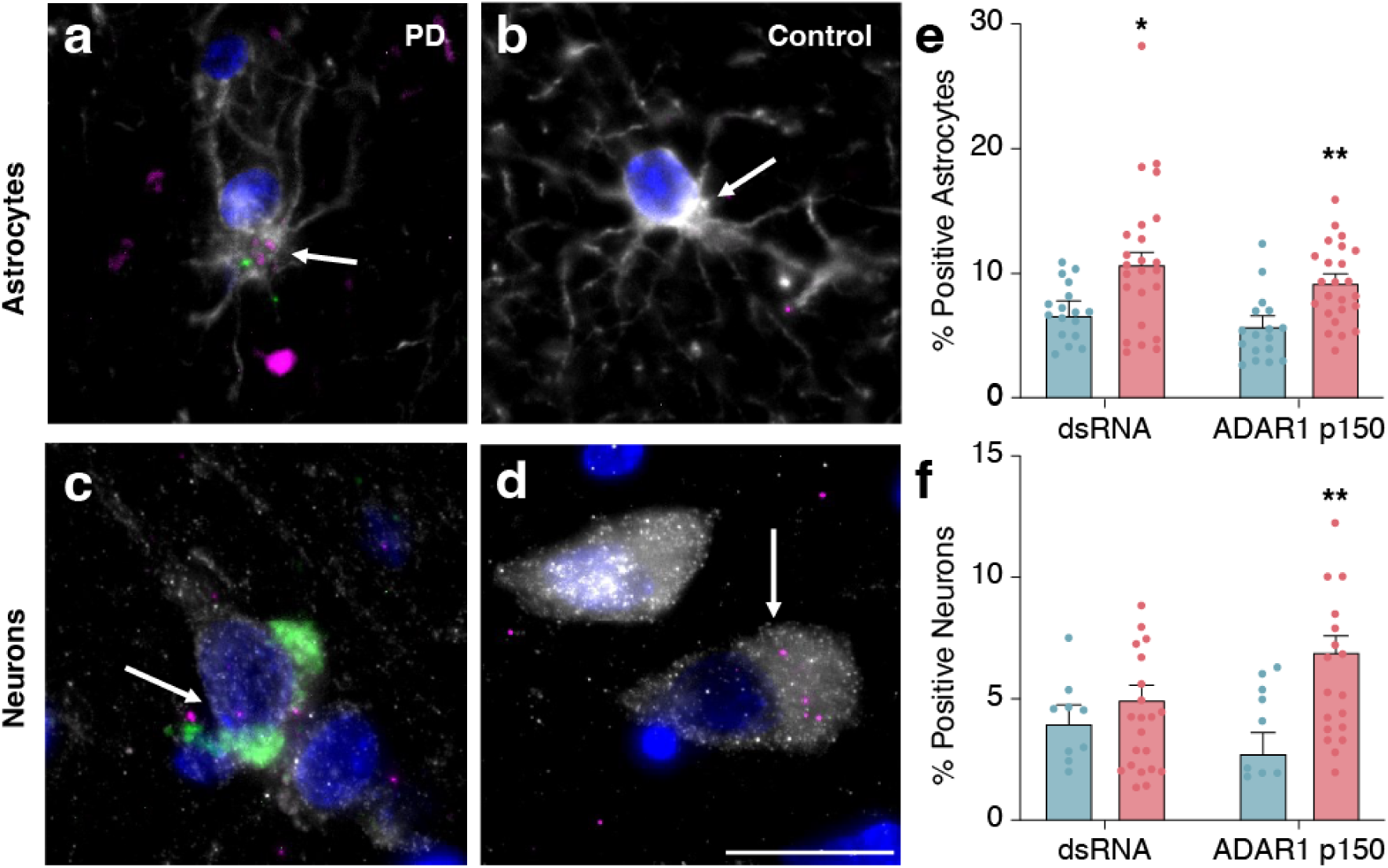
dsRNA accumulates selectively in astrocytes, and ADAR1 p150 increases in both astrocytes and neurons, in human Parkinson’s disease (PD) cortex. Representative immunofluorescence images of cortical astrocytes (GFAP (grey) and DAPI (blue) in a, b) and neurons (MAP2 (grey) and DAPI (blue) in c, d) also labelled for dsRNA J2 (green) and ADAR1 p150 (magenta) in PD (a, c) and control (b, d) cases. Arrows indicate ADAR1 p150-positive puncta expression. Scale bar = 20 µm. Quantification of the proportion of GFAP-positive astrocytes (e) and MAP2-positive neurons (f) positive for dsRNA J2 and ADAR1 p150 in the control and PD groups. Statistical analyses were conducted using SPSS. Data are presented as mean ± SEM with individual case data points. *P < 0.05, **P < 0.01.

In the cortical region examined, phosphorylated α-synuclein was detected in both astrocytes and neurons in PD cases (Fig. 3a,d) with a similar proportion of positive cells in the two cell types (3.27 ± 0.56% and 3.08 ± 0.46%, respectively, Fig. 3b,e). Triple immunofluorescence for ADAR1 p150, dsRNA and phosphorylated α-synuclein identified a higher frequency of triple-positive astrocytes in PD cortex than in controls (0.71 ± 0.09% vs 0.27 ± 0.11%, *P* = 0.006, Fig. 3c) and a modest positive correlation with astrocytic α-synuclein burden (Spearman ρ = 0.41, *P* = 0.007, *n* = 41). In contrast, the proportion of triple immunopositive neurons did not differ significantly between PD and controls (0.12 ± 0.04% vs 0%, *P* = 0.074, Fig. 3f). Together, these findings indicate that the association between dsRNA, ADAR1 p150 accumulation and α-synuclein pathology is more prominent in astrocytes than in neurons.

**Figure 3.**
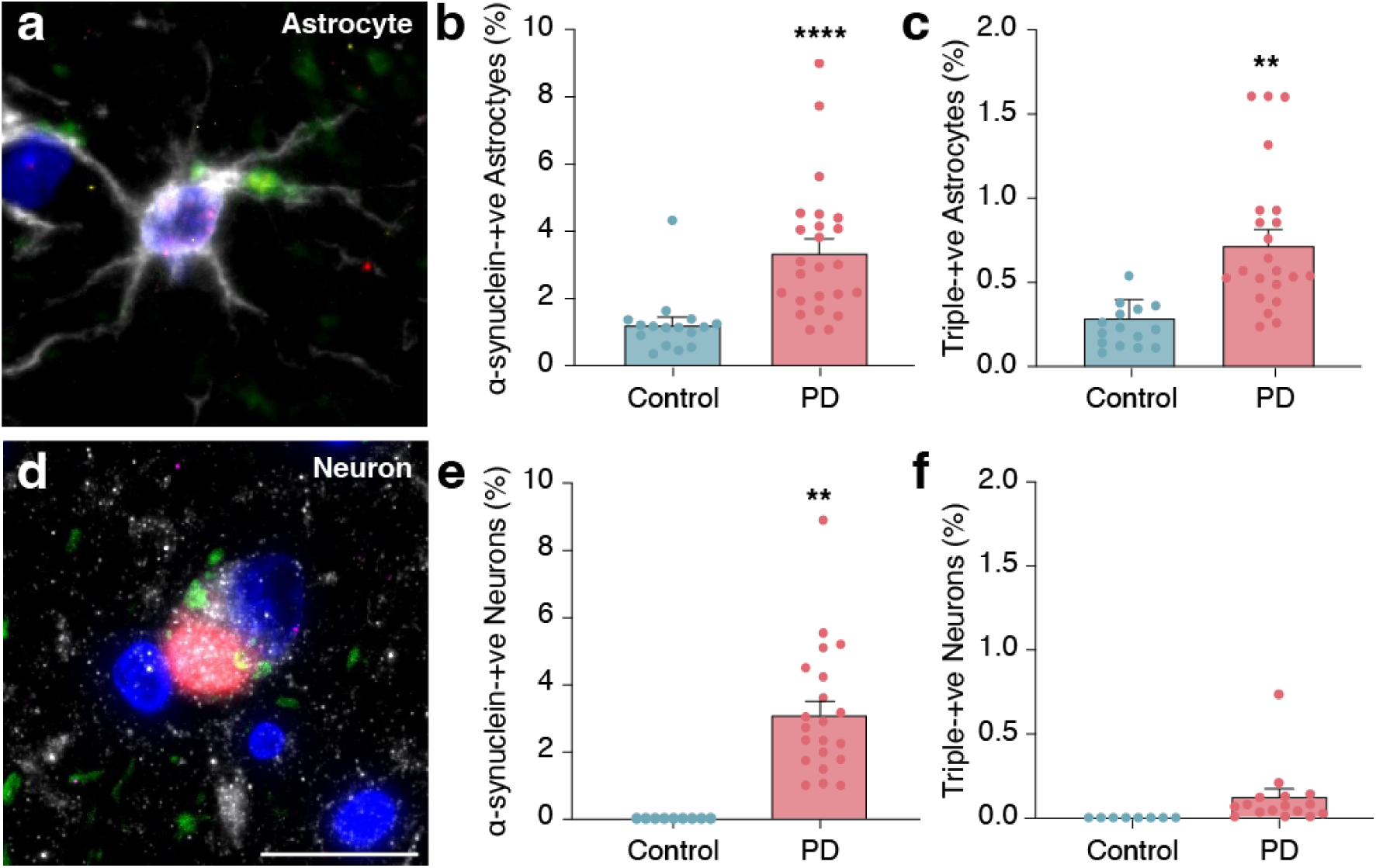
Astrocytic specific co-expression of α-synuclein, ADAR1 p150 and dsRNA in the human PD cortex. Immunofluorescence images of PD cortex stained for dsRNA/J2 (green), ADAR1 p150 (magenta), α-synuclein (red) and DAPI (blue) in astrocytes (a, GFAP (grey)) and neurons (d, MAP2 (grey)). Scale bar = 20 µm. Quantification of the proportion of α-synuclein-positive in astrocytes (b) and neurons (e). Quantification of the proportion of triple-positive (α-synuclein, ADAR1 p150 and dsRNA J2) among astrocytes (c) and neurons (f) in controls and PD cases. Data are presented as mean ± SEM with individual case data points. Statistical analyses were conducted using SPSS. \*\**p* < 0.01, \*\*\*\**p* < 0.0001.

We also examined whether the astrocytic dsRNA–ADAR1 phenotype is present in genetically driven α-synuclein cases (n = 3 *SNCA* mutation carriers) and show that similar selective astrocytic dsRNA, ADAR1 p150 and α-synuclein immunoreactivity occurs in such cases.

### Cell-type-specific transcriptomics of the dsRNA–ADAR1 pathway

To extend the protein-level findings, we analysed snRNA-seq data from 43 cortex donors using an eight-module gene scoring framework for astrocytes and total neurons in the cortex. Donor matched comparisons revealed clear cell-type partitioning of the dsRNA-related pathways (Fig. 4). Differences are expressed as Δ, the astrocyte-minus-neuron difference in mean module score on the log-normalised expression scale, such that positive values indicate higher scores in astrocytes and negative values indicate higher scores in neurons. Neurons showed significantly higher scores of the dsRNA editing and processing machinery: ADAR1 (Δ = −0.40, *P_adj* = 7.5 × 10^-9^, Fig. 4a), ADAR2 (Δ = −1.25, *P_adj* = 4.2 × 10^-11^, Fig. 4b), SNCA (Δ = −1.76, *P_adj* = 1.8 × 10^-12^, Fig. 4d), and the dsRNA processing module (Δ = −0.24, *P_adj* = 1.7 × 10^-9^, Fig. 4c). Conversely, astrocytes showed significantly higher scores for dsRNA sensing (Δ = 0.40, *P_adj* = 3.6 × 10^-9^, Fig. 4f), innate immune response (Δ = 0.15, *P_adj* = 7.5 × 10^-3^, Fig. 4e), and TE control (Δ = 0.39, *P_adj* = 2.7 × 10^-12^, Fig. 4h). The dsRNA/EV transport module did not differ significantly between the two cell types (Δ = 0.02, *P_adj* = 0.45, Fig. 4g). We note that these absolute-level comparisons describe baseline expression rather than functional engagement. The functional question of whether dsRNA pathway components are co-ordinately regulated with downstream immune programmes is addressed at the coupling level in subsequent sections.

**Figure 4.**
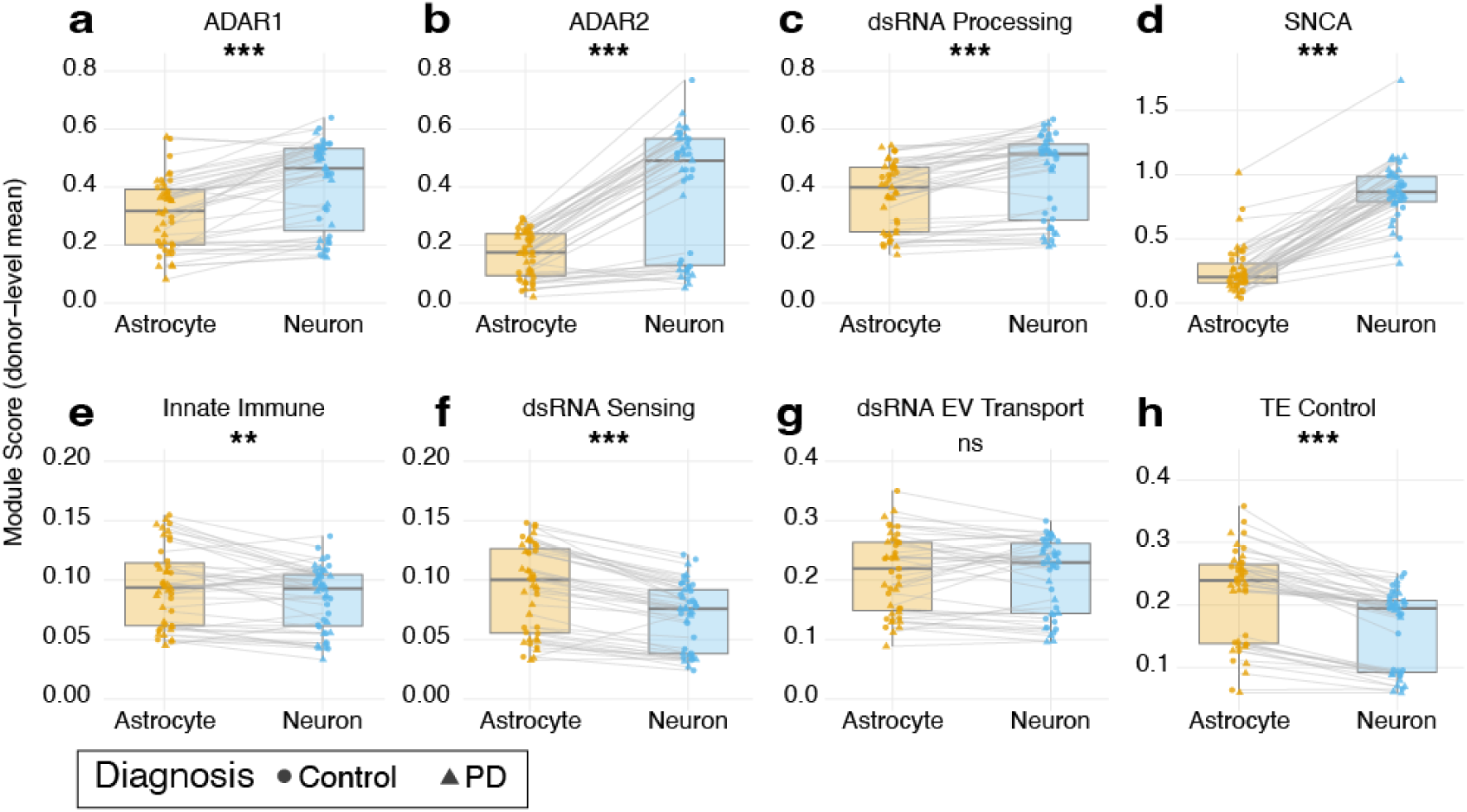
Cell type specific partitioning of dsRNA biology, editing machinery, and innate immune modules in the human cortex. Donor-level module scores are shown for astrocytes and total neurons from 43 donors (22 PD and 21 controls) across GSE178146, GSE202210 and GSE253462. Panels show the ADAR1 (a), ADAR2 (b), dsRNA processing (c), SNCA (d), innate immune response (e), dsRNA sensing (f), dsRNA EV transport (g), and TE control (h) modules. Each panel shows paired donor scores in astrocytes (yellow) and total neurons (blue); thin grey lines connect the same donor across the two cell types. Boxplots show the median and interquartile range. astrocyte versus neuron scores were compared within donor using two-sided paired Wilcoxon signed-rank tests, with BH correction across the eight modules: ** *P_adj* < 0.01, *** *P_adj* < 0.001, ns = not significant.

No significant PD versus control differences were detected in any of the eight module scores in any cell type after covariate adjustment and BH correction (all *P_adj* > 0.29; Supplementary Information Fig. 2). Consistent with the module-level analysis, pseudobulk differential-expression analysis using DESeq2 identified no genes within the eight modules that reached significance in astrocytes or total neurons after BH correction.

Among the 31 donors with available Braak staging (stage 0, n=15, stages 4–5, n=5, stage 6, n=11), no module showed a Braak-dependent change in either astrocytes or total neurons after false discovery rate (FDR) correction, and none replicated across the two cohorts (Supplementary Information Fig. 3a). Within-donor astrocyte–neuron divergence was likewise flat: no module reached FDR significance in the primary astrocyte vs total neuron contrast, with sensitivity analyses using excitatory or inhibitory neurons as the reference yielding the same null pattern (Supplementary Information Fig. 3b).

### Complement shows the strongest positive and most cell-type-divergent coupling with dsRNA sensing in astrocytes

To determine whether the cell-type differences in dsRNA-related module scores were associated with distinct downstream immune programmes, we calculated donor-level partial Spearman correlations between each of the seven dsRNA modules and nine innate immune sub-panels in astrocytes and total neurons (n = 43). Comparison of the correlation matrices using the paired donor-level bootstrap revealed extensive cell-type divergence (Fig. 5a,b). Complement showed the most robust astrocyte-dominant pattern. In astrocytes, complement scores were positively correlated with dsRNA-related module scores (ρ = 0.36 to 0.58, Fig. 5a), whereas in neurons, complement showed uniformly negative correlations at six of the seven modules (ρ = −0.53 to −0.70, Fig. 5b); the association with the SNCA module was not significant (*P_adj* > 0.05). The TLR-DAMP sub-panel also showed significant astrocyte-specific coupling for six of the modules (all *P_adj* < 0.003) with no evidence of coupling for the SNCA module (*P_adj* = 0.89) and TLR-dsRNA reached significance for the dsRNA sensing module (*P_adj* = 0.016). Notably, dsRNA sensing was negatively correlated with the astrocyte reactivity sub-panel in astrocytes (ρ = -0.43, *P_adj* = 0.01), indicating that the positive dsRNA sensing–complement relationship was not accompanied by a general increase across all astrocyte-reactivity programmes. For the remaining sub-panels the correlation with dsRNA sensing had the same sign in both cell types and differed only in magnitude: inflammasome (astrocytes, ρ = −0.55, neurons ρ = −0.74), NF-κB (astrocytes, ρ = −0.42, neurons ρ = −0.27), ISG core (astrocytes, ρ = −0.42, neurons ρ = −0.50), TLR-dsRNA (astrocytes, ρ = 0.40, neurons, ρ = 0.11) and OAS-RNase L (astrocytes, ρ = −0.72, neurons ρ = −0.85). Apart from complement and the TLR sub-panels, all of these canonical dsRNA-response readouts including ISG core, OAS–RNase L and NF-κB and inflammasome correlated negatively with the dsRNA sensing score in astrocytes. Complement is therefore not simply the strongest member of a coherent, uniformly positive antiviral response; it is the only effector programme in the panel whose donor-level covariation with dsRNA sensing is positive in astrocytes.

**Figure 5.**
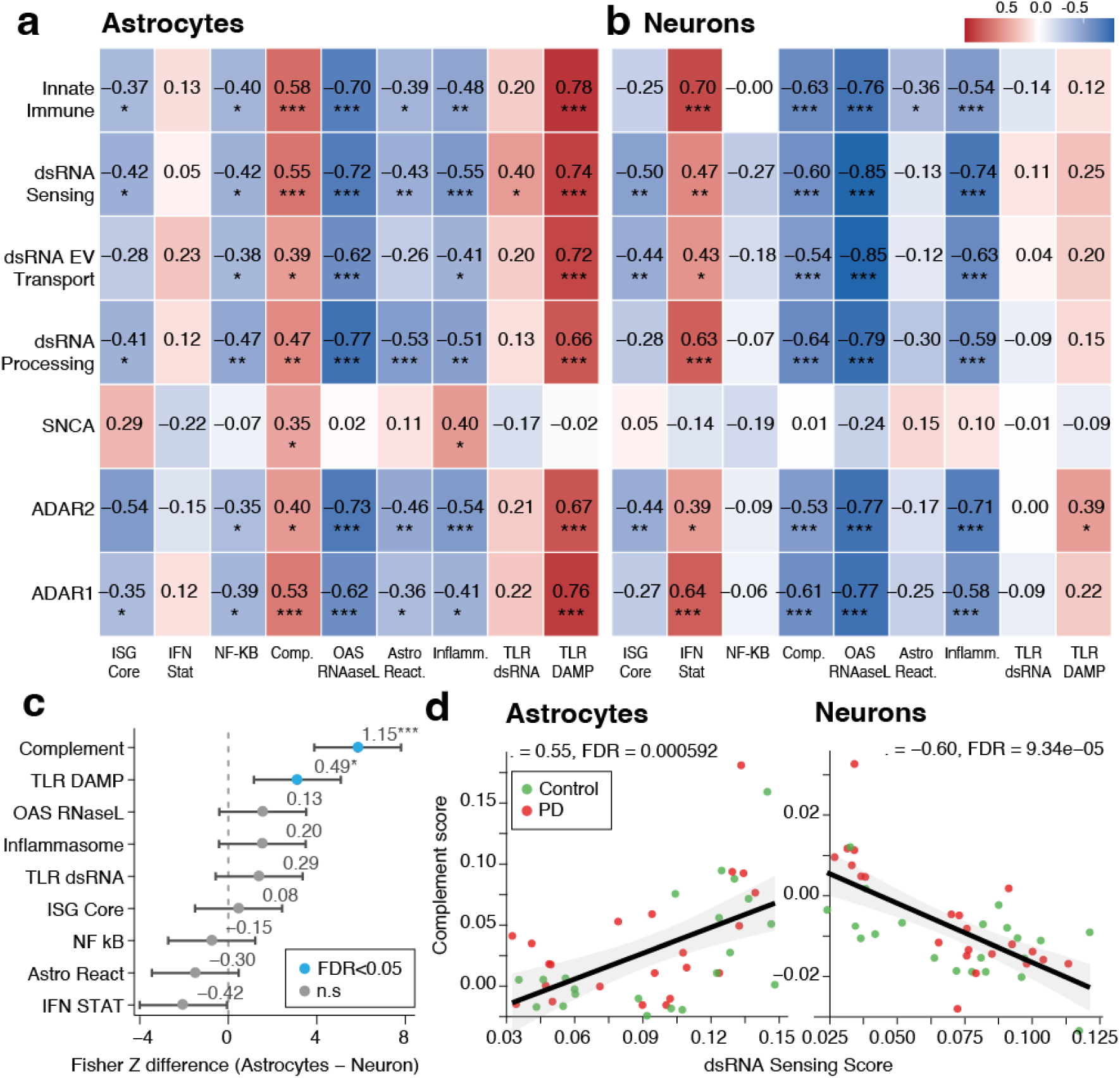
dsRNA sensing shows cell-type-divergent coupling with the complement programme in the human cortex. (a,b) Heatmap of partial Spearman correlations for astrocytes (a) and total neurons (b) between seven dsRNA-related modules and nine immune sub-panel scores. (c) Forest plot of the astrocytes versus neurons difference in the coupling of dsRNA sensing with each of the nine immune subpanels. Points and horizontal bars show paired donor-bootstrap estimates and 95% confidence intervals; blue indicates FDR < 0.05 and grey indicates non-significant. (d) Scatter plot showing the association between complement and dsRNA sensing scores in astrocytes (left) and total neurons (right), stratified by diagnostic group (red, PD; green, control). Module and panel scores were background-subtracted (panel mean minus the mean of expression-bin-matched control genes). Group-specific Spearman’s ρ values and significance are shown within each plot, * P_adj < 0.05, ** P_adj < 0.01, *** P_adj < 0.001.

We next quantified the astrocyte-versus-neuron difference in coupling between the dsRNA sensing and each of the immune sub-panels. Complement showed the largest difference between cell types (ρ_astrocyte = 0.55, ρ_neuron = -0.60 across all 43 donors; Δρ = 1.15, *P_adj* < 0.0001, Fig. 5c), and the positive association in astrocytes was present in both diagnostic groups, in controls (ρ = 0.70) and in PD (ρ = 0.53), with no significant difference between them (*P_adj* = 0.71, Fig. 5d). In neurons, the corresponding correlations were consistently negative in both PD cases (ρ = −0.77) and controls (ρ = −0.50). Thus, the direction of the dsRNA sensing-complement relationship differed consistently between astrocytes and neurons, and was stable across disease groups, rather than emerging only in disease.

TLR-DAMP showed a smaller but significant difference in the same direction (Δρ = 0.49, *P_adj* < 0.01), whereas IFN-STAT diverged in the opposite direction, reflecting stronger coupling in neurons, although this comparison did not remain significant after multiple testing correction (Δρ = −0.42, *P_adj* > 0.05). The remaining sub-panels, including inflammasome, NF-κB, ISG core, TLR-dsRNA and OAS-RNase L, showed no significant astrocyte-neuron differences in their correlations with dsRNA sensing. These results identify complement as the immune programme showing the strongest cell-type-divergent relationship with dsRNA sensing.

### Transposable element (TE) control is tightly associated with dsRNA sensing and innate immune modules

The TE control module captures genes involved in the epigenetic and post-transcriptional suppression or processing of TE (48–50, 53). Because failure or compensatory engagement of TE-control mechanisms may be associated with endogenous dsRNA accumulation, we examined whether variation in this module was associated with other components of the dsRNA-related programme. Importantly, the TE-control module measures expression of TE-control machinery and does not directly quantify TE expression, TE derepression or TE-derived dsRNA. Across 43 donors, donor-level partial Spearman correlation analysis showed a strong positive association between TE control and multiple dsRNA-related modules in both astrocytes and neurons (Fig. 6a). In astrocytes, TE control score was associated with the innate immune module (ρ = 0.94), dsRNA processing (ρ = 0.92), ADAR1 (ρ = 0.91), dsRNA sensing (ρ = 0.89) and ADAR2 (ρ = 0.79) modules (all *P_adj* < 1 × 10^-8^). A similar pattern was observed in total neurons, TE control score was strongly associated with innate immunity (ρ = 0.94), dsRNA processing (ρ = 0.97), and with ADAR1 (ρ = 0.93, all *P_adj* < 10^-5^). Subdivision of neurons into excitatory and inhibitory populations did not alter this pattern, with correlations ranging from ρ = 0.83 to 0.97 across both subsets. By contrast, SNCA was the only module not significantly correlated with TE control in either cell types (astrocytes: ρ = -0.09, *P_adj* = 0.56; total neurons: ρ = 0.13, *P_adj* = 0.41). Importantly, TE control score was also positively associated with the complement sub-panel in astrocytes (ρ = 0.50, *P_adj* = 2.08 × 10□³, Fig. 6b), and negatively associated with complement in neurons (ρ = -0.70, *P_adj* = 1.68 × 10^-6^). This bidirectional divergence further supports the link between endogenous dsRNA-related pathways and astrocyte-selective complement related gene expression. The correlations among the dsRNA-related modules themselves are uniformly high in both cell types (ρ = 0.79–0.97). Correlations of this magnitude between nominally distinct gene sets are unlikely to reflect pathway-specific biology alone and probably also capture a shared donor-level axis of global transcriptional output and detection efficiency. The interpretable signal is the reversal in sign of the complement association between astrocytes and neurons, which a shared technical axis cannot generate, rather than the absolute magnitude of the module–module correlations.

**Figure 6.**
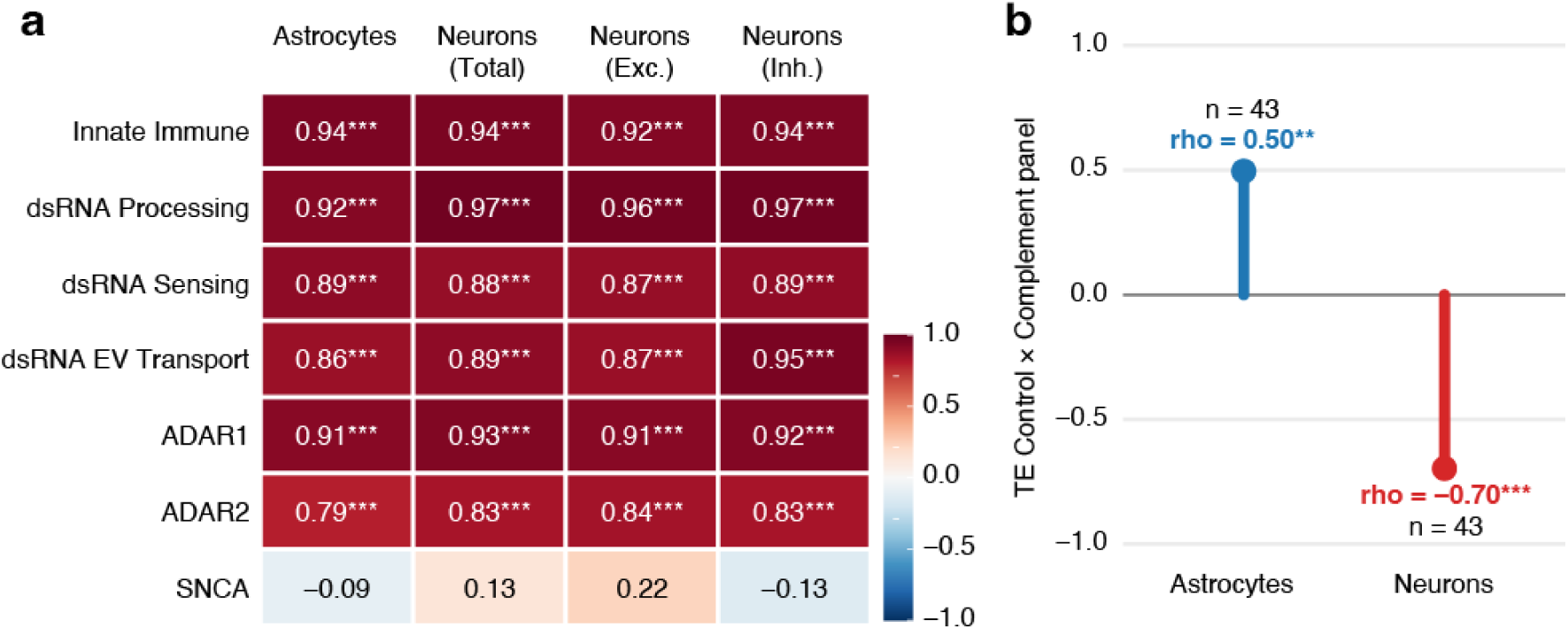
Strong associations between the TE control module and dsRNA-related modules, with cell-type-divergent associations with complement in the cortex. (a) Heatmap of donor level correlation between the TE control module score and the seven other functional modules in astrocytes and neurons. Colour scale indicates correlation coefficients from −1 to +1. (b) Lollipop plot of correlation between TE control and the complement sub-panel in the astrocytes (blue) and neurons (red). Lollipop length indicates the magnitude of the correlation coefficients. Significance is based on BH-corrected test: *** P_adj < 0.001.

### Reactive astrocyte gene signatures show broad associations with dsRNA–ADAR1-innate immune axis

To determine whether astrocytic dsRNA sensing and complement coupling was associated with particular reactive astrocyte signatures, we calculated five composite scores from donor-level astrocyte pseudobulk profiles: A1-like all, A1-like complement-enriched, A1-like non-complement, A2-like and pan-reactive signatures. Each score was correlated with the seven prespecified functional modules across all 43 donors (Fig. 7a).

**Figure 7.**
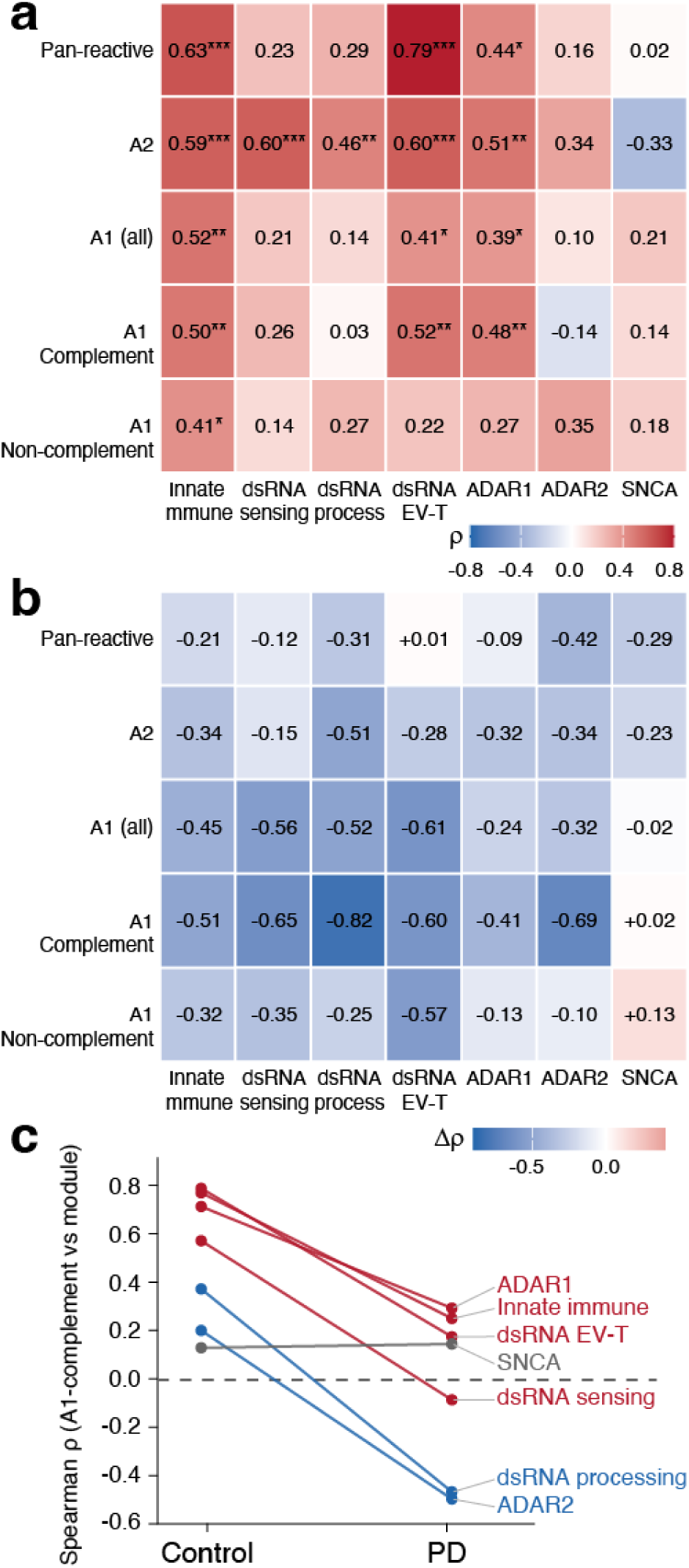
Reactive-astrocyte gene signature associated with the dsRNA-ADAR1-innate-immune axis modules. (a) Donor-level pseudobulk Spearman correlations between five astrocyte subgroup composite scores (rows) and seven astrocyte module scores (columns). (b) Descriptive difference in correlation between PD and control donors, calculated as Δρ = ρ_PD − ρ_Control. (c) Control to PD slope plot for the correlations of A1-like complement enriched signature. dsRNA EV-T = dsRNA EV-Transport. Cell values indicate ρ; significance is based on BH-corrected test: *P_adj < 0.05, ** P_adj < 0.01, *** P_adj < 0.001.

Across all donors, the innate immune module was positively correlated with all five reactive astrocyte composites (ρ = 0.40-0.63, all *P_adj* < 0.023), and the dsRNA EV transport module was correlated with all composites except A1-like non-complement (ρ = 0.41-0.79). The A2-like composites showed the broadest overall pattern, correlating significantly with five of seven modules (ρ = 0.46 to 0.60). Within the A1-related signatures, the A1-complement composite, built from *CFB, C1R, C1S, C3, CFI, SERPING1*, was positively correlated with dsRNA EV transport (ρ = 0.52, *P_adj* = 2.2 × 10 ³), the innate immunity module (ρ = 0.51, *P_adj* = 2.9 × 10 ³) and ADAR1 (ρ = 0.48, *P_adj* = 5.4 × 10 ³), whereas A1-non-complement composite (*FKBP5, GBP2, PSMB8*) was significantly correlated with innate immune module alone (ρ = 0.41, *P_adj* = 0.022) and was not significantly correlated with ADAR1 (ρ = 0.27, *P_adj* = 0.15). SNCA did not show a statistically significant association with any of the five reactive-state composites (|ρ| ≤ 0.33, all *P_adj* ≥ 0.077), consistent with its relative independence from the other dsRNA-related modules.

Diagnostic-group stratification suggested that several associations between the A1-like complement-enriched signature and dsRNA-axis modules were numerically stronger in controls than in PD cases (Fig. 7b,c). In controls, this signature was positively correlated with dsRNA EV transport (ρ = 0.78), innate immunity (ρ = 0.76), ADAR1 (ρ = 0.70) and dsRNA sensing (ρ = 0.56, all *P _adj* < 0.027). In PD, the corresponding point estimates were lower and ranged from -0.08 to 0.29, and none reached significance. However, formal bootstrap comparison of the control and PD correlations showed that none of these group differences reached significance after multiplicity correction. SNCA coupling was comparable across groups (ρ = 0.13–0.15) and did not display the same numerical group pattern. Accordingly, the wide confidence intervals and non-significant group-comparison tests do not provide evidence that coupling is attenuated in PD.

Together, the all-donor analysis demonstrates that several historical reactive-astrocyte gene signatures covary with the dsRNA–ADAR1–innate immune axis. The A2-like signature showed the broadest overall association pattern, whereas the A1-like complement-enriched signature was specifically associated with ADAR1, dsRNA EV transport and innate immune activity. These donor-level composite relationships indicate coordinated transcriptional variation but do not establish the presence of discrete A1 or A2 astrocyte populations. The lower correlation estimates observed in PD for the A1-like complement-enriched signature remain exploratory and require validation in larger independent cohorts.

### ADAR1-ranked gene programmes are enriched for neurodegeneration-associated pathways

Gene set enrichment analysis (fGSEA with KEGG) was performed across all eight modules, ranking genes by their covariate-residualised Spearman correlation with each module score. Both cell types showed extensive enrichment in the cortex (unique pathways significant in at least one of the eight module analyses; astrocytes: 95 significant pathways; neurons: 92; *P_adj* < 0.05, Supplementary Datasheets), indicating that the dsRNA–ADAR pathway is embedded within broad cellular programmes in both cell types.

To identify the functional programmes associated with ADAR1 in astrocytes, we focused on genes ranked by their correlation with the ADAR1 module score. In astrocytes, 50 KEGG pathways were significantly positively enriched (*P_adj* < 0.05). Oxidative phosphorylation showed the strongest enrichment (NES = 2.26, *P_adj* = 1.4 × 10^-10^; Fig. 8a,c), followed by the PD pathway (NES = 2.17, *P_adj* = 2.5 × 10^-9^), Huntington’s disease (NES = 2.04, *P_adj* = 4.9 × 10^-10^), and Alzheimer’s disease (NES = 1.98, *P_adj* = 3.0 × 10^-8^). These findings associate the astrocytic ADAR1-ranked gene programme with mitochondrial neurodegeneration-related pathways.

**Figure 8.**
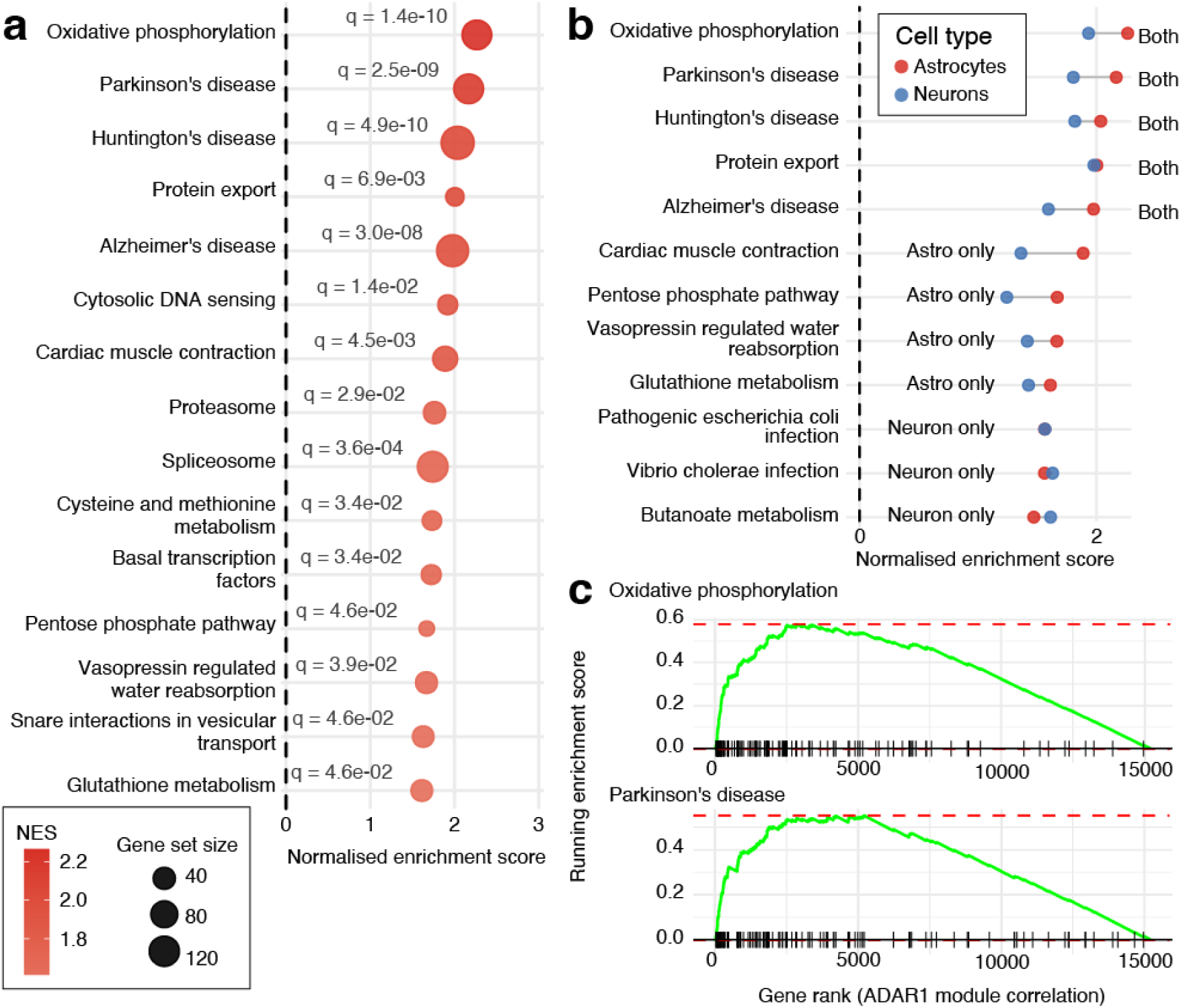
ADAR1-correlated pathway enrichment in astrocytes. (a) Dot plot showing the top 15 significantly enriched KEGG pathways (P_adj < 0.05) from fGSEA of ADAR1-correlated gene rankings in astrocytes. Dot size represents gene set size; colour represents NES. Oxidative phosphorylation (NES = 2.26) and PD (NES = 2.17) are the top-ranked pathways. (b) Comparison of ADAR1-correlated pathway enrichments between astrocytes and neurons for selected pathways. (c) fGSEA enrichment plots for oxidative phosphorylation and PD pathways.

Additional ADAR1-correlated pathways in astrocytes included proteasome (NES = 1.76, *P_adj* = 0.029) and cytosolic DNA sensing (NES = 1.92, *P_adj* = 0.014). Beyond the ADAR1 module, JAK-STAT signalling reached significance in several other astrocyte module analyses (NES = 1.81–1.96, *P_adj* < 10^-3^).

Comparable mitochondrial and proteostasis pathway enrichments were also detected in ADAR1-ranked neuronal gene programmes (52 significant positively enriched pathways for the ADAR1 module). Among pathways significant in both cell classes, the descriptive NES values were higher in astrocytes, led by oxidative phosphorylation (astrocytes NES = 2.26 vs neurons NES = 1.94, Fig 8b), Parkinson’s disease (2.17 vs 1.81), Huntington’s disease (2.04 vs 1.82) and protein export (2.00 vs 1.98). Cytosolic DNA sensing was also significant in both, though with a larger astrocytic effect (1.92 vs 1.60). These results indicate that the ADAR1–mitochondrial coupling is not exclusive to astrocytes at the transcriptomic level. The astrocyte-specific interpretation is anchored by the immunofluorescence findings, which demonstrate that dsRNA and ADAR1 p150 protein accumulate preferentially in astrocytes in PD cortex (Fig. 2).

Astrocyte-leading enrichments included glutathione metabolism (astrocytes NES = 1.61 vs neurons NES = 1.43, Fig 8b), the pentose phosphate pathway (1.67 vs 1.24) and vasopressin-regulated water reabsorption (1.67 vs 1.42), the last of these being driven by aquaporin-family and vesicular trafficking components. Several pathways met the significance threshold only in the neuronal ADAR1-ranked analysis, including ubiquitin-mediated proteolysis (NES = 2.25), endocytosis (NES = 1.84), autophagy regulation (NES = 1.85) and mTOR signalling (NES = 1.69, Supplementary Datasheets). These findings highlight proteostatic and trafficking programmes associated with ADAR1-ranked genes in neurons.

## Discussion

### A cell-type-specific division of labour in the dsRNA pathway

In this study, we provide an integrated protein- and transcriptomic-level characterisation of the dsRNA–ADAR1 axis across cell types in the human cortex. Three convergent lines of evidence position astrocytes as the primary cellular site of dsRNA pathway activity. Immunofluorescence demonstrates a PD-associated increase in dsRNA in GFAP-positive astrocytes together with ADAR1 p150 induction in both astrocytes and neurons, and astrocytic ADAR1 p150 correlating with α-synuclein pathology; transcriptomics revealed astrocyte enrichment for the dsRNA sensing, innate immune, and TE control modules, and at the network level, complement activation emerged as the predominant astrocyte-specific effector programme coupled to dsRNA activity, whereas the coupling of dsRNA sensing to NF-κB and ISG programmes did not differ significantly between cell types. Critically, this co-regulation was present in controls as well as PD and did not differ significantly between diagnostic groups, indicating a constitutive feature rather than an activation event induced by disease. Point estimates of coupling were numerically lower in PD across the disease-stratified contrasts, but no group difference reached significance and the confidence intervals were wide. We therefore treat destabilisation of this axis in disease as a hypothesis to be tested in larger samples.

The cell-type partitioning with neurons expressing higher levels of ADAR1, ADAR2, and SNCA while astrocytes are enriched for dsRNA sensing and innate immunity, parallels recent transcriptomic findings. Multiple human PD snRNA-seq studies have reported a consistent pattern of transcriptional repression in neurons alongside glial activation, with astrocytes showing reactive and neuroinflammatory phenotypes (54, 55). Ma *et al.* documented neuroinflammatory pathway activation in PD astrocytes, including JAK/STAT and NF-κB signalling, alongside a CD44-positive reactive astrocyte state that gained spatial correlation with infiltrating T cells (34). Feleke *et al.* reported that in the anterior cingulate cortex, Lewy body diseases were characterised by transcriptional alterations across multiple cell types, with excitatory neurons and oligodendrocytes most prominently affected, and widespread RNA splicing dysregulation particularly in the dementia patients (33). Our findings extend this body of work by identifying a specific molecular axis of the dsRNA–ADAR1 pathway that partitions cleanly across cell types.

In our data, neurons express higher absolute ADAR1 mRNA than astrocytes, yet astrocytes show the dsRNA-coupled innate immune signature both at the protein level (Figs. 2, 3) and the network level (Fig. 5). The constitutive ADAR1 p110 isoform is predominantly nuclear and edits intronic and other nuclear dsRNA that are largely irrelevant to cytosolic MDA5/RIG-I sensing, while the interferon-inducible ADAR1 p150 isoform is cytoplasmic and is the editor whose loss permits inappropriate MDA5 activation by endogenous dsRNA (22, 56, 57). The p110 and p150 isoforms differ at their 5’ ends but share their 3’ ends (58). 3’-biased 10X snRNA-seq cannot distinguish between them, and total ADAR1 mRNA in our dataset may therefore report the abundant constitutive neuronal p110 pool, which is not expected to track inflammatory pathways (56). The higher neuronal score on editing and processing modules is consistent with the high constitutive A-to-I editing load of neurons. However, the canonical recoding events at synaptic transcripts are ADAR2-dependent, so this neuronal signal should not be attributed to ADAR1 p110 alone (56). By contrast, our immunofluorescence is p150-specific and demonstrates preferential p150 accumulation in astrocytes that correlates with α-synuclein burden, consistent with prior reports that α-synuclein oligomers trigger ADAR1 isoform switching toward p150 in iPSC-derived astrocytes (25). Within this framework, the astrocyte-specific coupling between ADAR1 module variation and downstream immune programmes is likely to reflect variation in the IFN-inducible p150 fraction rather than in total ADAR1 levels, linking the absolute-expression and coupling-architecture layers into a single coherent picture.

Garza *et al.* (19) demonstrated that the majority of upregulated TEs in PD brain were endogenous retroviruses (ERV) from the ERV-K, ERV-W, and ERV-H subfamilies, many retaining near-intact proviral structures. Their observation that TE activation and interferon responses were restricted to the nigrostriatal pathway, with no TE upregulation in the prefrontal cortex, is consistent with the absence of a detectable PD-versus-control transcriptomic difference in our cortical DESeq2 analysis. The cortex may maintain a well-calibrated editing and sensing equilibrium that prevents TE-driven inflammation, whereas the nigrostriatal system, under greater oxidative stress, may exceed this buffering capacity. The donor-stable coupling architecture we describe in cortical astrocytes is consistent with such regional buffering and offers a testable framework for why cortical involvement in PD lags that in the nigrostriatal pathway.

### Complement as an astrocyte-specific effector of dsRNA-driven innate immunity

The central transcriptomic finding is that complement is the predominant innate immune programme showing astrocyte-specific coupling with dsRNA pathway activity correlating with astrocytic α-synuclein accumulation. Complement sub-panel scores positively correlated with dsRNA modules in astrocytes (ρ = +0.35 to +0.58) and negatively in neurons (ρ = −0.53 to −0.70), a finding that was independent of PD. In fact, point estimates of coupling were numerically higher in controls than in PD, the opposite of what would be expected if PD newly induced the pathway. This bidirectional pattern suggests not merely cell-type-restricted expression but opposing co-regulatory architectures, possibly reflecting the known role of astrocytes as the primary CNS source of complement (51, 59) with neuronal programmes actively suppressing complement tagging (60). Because our IF data show that the substrate for this circuit, astrocytic dsRNA and ADAR1 p150, increases in PD, the most parsimonious interpretation is that PD raises the input to a pre-existing complement-competent circuit rather than one that activates a new inflammatory pathway. This may be due to astrocytic accumulation of α-synuclein perturbing this axis through transcriptional derepression of repetitive elements, sequestration of RNA-binding proteins, and impaired RNA quality control (25, 61).

The cell-type divergence pattern of the response requires explicit comment, as within astrocytes the dsRNA sensing score also correlated negatively with the ISG core, OAS–RNase L, NF-κB and inflammasome sub-panels. This is not the profile expected if higher constitutive sensing capacity simply produced a proportionally stronger canonical antiviral response, and it constrains how the complement result should be interpreted. One interpretation is that the response is homeostatic: donors whose astrocytes maintain a high sensing and editing apparatus may be those in whom endogenous dsRNA is efficiently contained, so that interferon-stimulated and inflammasome output remains low, while complement, which in astrocytes is regulated substantially through NF-κB-independent and developmentally constitutive routes, tracks the surveillance machinery itself rather than its activation state. An alternative interpretation is technical: interferon-stimulated transcripts are expressed at low levels in postmortem cortex and are sensitive to agonal and postmortem factors, so donor-level ISG scores may be dominated by variance unrelated to dsRNA burden. We cannot adjudicate between these possibilities and therefore describe complement as the sub-panel whose covariation with dsRNA sensing is both positive and cell-type-divergent, without inferring that dsRNA sensing activates a canonical interferon programme in these astrocytes. The IFN-STAT coupling with the innate immune module (*P_adj* = 0.010) further suggests that the downstream routing of interferon signals differs between cell types, consistent with astrocytes as the principal CNS source of complement components (51, 62) and neurons engaging IFN signalling for distinct, non-complement effector programmes (63, 64).

Our findings support recent reports identifying that α-synuclein activates astrocytic complement secretion via NF-κB (65), the same pathway engaged by cytoplasmic dsRNA through MDA5/MAVS signalling (66). Importantly, dsRNA sensing was negatively correlated with the astrocyte reactivity sub-panel. This argues against the alternative explanation that the complement coupling is simply a proxy for general reactive gliosis, dissociating complement co-regulation from the broad pan-reactive astrocyte response. In PD cortex there is a change of input to a pre-existing RNA-surveillance circuit rather than the activation of a new one. The dsRNA EV transport module did not differ between astrocytes and neurons, indicating comparable baseline expression of the underlying membrane-trafficking machinery, yet it is among the strongest correlates of reactive-astrocyte composites. Because this module is dominated by generic vesicular and endolysosomal genes, the most parsimonious reading is coordinated upregulation of the secretory apparatus that accompanies astrocyte reactivity, rather than evidence for RNA transfer between cells, which these data cannot address.

### ADAR1 as a transcriptional hub linking RNA editing to mitochondrial and proteostasis stress

In astrocytes, ADAR1-correlated gene programmes converge on KEGG oxidative phosphorylation, the PD pathway, Huntington’s disease and Alzheimer’s disease gene sets, placing ADAR1 at a crossroads of mitochondrial metabolism, proteostasis and innate immunity. Comparable mitochondrial and proteostasis enrichments are also detected in neuronal ADAR1-ranked programmes (52 significant pathways), indicating that ADAR1 activity is not exclusive to astrocytes but reflects shared underlying biology in which ADAR1 sits at the interface of mitochondrial metabolism, proteostasis, and immunity across cell types. The cell-type-specific contextualisation of this signature in PD cortex is anchored by our immunofluorescence data, which show that the PD-associated dsRNA increase is confined to astrocytes, and ADAR1 p150 protein accumulation positivity correlated with α-synuclein burden.

Mitochondrial dysfunction is a well-established feature of PD astrocytes (67–69). The cytosolic DNA sensing pathway enrichment is of particular interest, as it suggests crosstalk between dsRNA and dsDNA sensing through shared downstream effectors such as TBK1 and STING (70). This is consistent with evidence that mitochondrial DNA release from damaged mitochondria activates cGAS–STING signalling, which converges with the MDA5–MAVS axis on type I IFN induction (71). JAK-STAT signalling reached significance across multiple modules in astrocytes, with higher enrichment scores than in neurons, consistent with the astrocyte-dominant immune coupling identified in the coupling analysis.

### Limitations

This study has several limitations. First, snRNA-seq cannot resolve ADAR1 p110 and ADAR1 p150 isoforms due to 3′-bias; isoform-resolved long-read RNA sequencing would provide more definitive evidence for p150-specific induction in astrocytes. Second, the immunofluorescence cohort (23 sporadic PD, 18 controls, 3 *SNCA* carriers) and the snRNA-seq cohort (22 PD, 21 controls) are independent samples, so the integration is convergent rather than donor-matched. Third, with 43 cortical donors (22 PD, 21 control) our differential-expression analysis is powered to detect only moderate-to-large PD versus control effects. The module-level and the coupling-level null results are therefore evidence against large PD-versus-control effects rather than proof of no effect. Fourth, standard 10X 3′ libraries have poor sensitivity for repetitive elements (19, 72), so our inference that TE-derived dsRNA engages the sensing/editing modules rests on correlative evidence and external validation rather than direct TE transcript measurement. Finally, cross-cell-type module correlations may retain residual confounding from shared library-size or biological variation despite covariate adjustment.

## Conclusions

This study integrated protein and transcriptomic characterisation of the dsRNA–ADAR1 axis across cell types in the PD brain. Immunofluorescence demonstrates preferential dsRNA accumulation in astrocytes and increased ADAR1 p150 protein in both astrocytes and neurons, with astrocytic ADAR1 p150 positively correlating with α-synuclein accumulation. Transcriptomic analysis of 43 cortical donors reveals a division of labour: neurons express the editing and processing machinery, astrocytes are enriched for dsRNA sensing, innate immunity, and TE control. Among nine innate immune sub-panels, complement activation shows the strongest and most consistent astrocyte-specific coupling with dsRNA pathway activity, while interferon- and NF-κB-related couplings are shared between cell types.

ADAR1-correlated programmes in astrocytes track mitochondrial and neurodegeneration-associated pathways, with weaker evidence for proteasomal gene sets. These findings reframe complement-mediated neuroinflammation in PD as the engagement of a constitutive, astrocyte-intrinsic RNA-surveillance circuit by a rising dsRNA burden, rather than the *de novo* induction of an immune programme. The dsRNA burden correlates with a-synuclein accumulation in astrocytes. This identifies dsRNA–ADAR1–complement coupling in astrocytes as a candidate mechanistic node for therapeutic targeting in PD.

## Supporting information

Supplementary Tables, Figures and References

Supplementary data sheets

## Declarations

## Ethics approval and consent to participate

The assessment of postmortem human brain tissue was approved by the Human Research Ethics Committee at the University of Sydney (2020/HE000707).

## Consent for publication

Not applicable.

## Data availability

The snRNA-seq datasets analysed in this study are publicly available in the Gene Expression Omnibus under accession numbers GSE202210, GSE178146, and GSE253462. All analyses are available at https://doi.org/10.5281/zenodo.21315999.

## Authors’ Contributions

YL performed experimental research and bioinformatic analysis, collected and analyzed data, and wrote the manuscript. YF and GH conceived and designed the study, edited the manuscript, and approved the final version.

## Acknowledgements

The authors acknowledge: human brain tissue sample procurement by the Sydney Brain Bank, Sydney (enabled by funding from Neuroscience Research Australia and is a special gift in memory of Jim Raftos from the Shaw family); Anahid Ansari Mahabadian and Amy Burke for research assistance; the use of Microscopy Australia infrastructure at the Sydney Microscopy and Microanalysis facility, University of Sydney (enabled by the National Collaborative Research Infrastructure Strategy). We thank Heidi Cartwright for assistance with figure preparation.

## List of abbreviations

A-to-I: adenosine-to-inosine
ADAR: adenosine deaminase acting on RNA
BH: Benjamini–Hochberg
CNS: central nervous system
DAMP: damage-/danger-associated molecular pattern
dsDNA: double-stranded DNA
ERV: endogenous retrovirus
EV: extracellular vesicle
FDR: false discovery rate
fGSEA: fast gene set enrichment analysis
GEO: Gene Expression Omnibus
GFAP: glial fibrillary acidic protein
IFN: interferon
iPSC: induced pluripotent stem cell
ISG: interferon-stimulated gene
JAK: Janus kinase
KEGG: Kyoto Encyclopedia of Genes and Genomes
MAP2: microtubule-associated protein 2
MAVS: mitochondrial antiviral signalling protein
MDA5: melanoma differentiation-associated protein 5 (encoded by IFIH1)
MSigDB: Molecular Signatures Database
NES: normalised enrichment score
NF-κB: nuclear factor κB
NLRP3: NLR family pyrin domain-containing 3
OXPHOS: oxidative phosphorylation
PD: Parkinson’s disease
QC: quality control
RIG-I: retinoic acid-inducible gene I (encoded by DDX58)
SEM: standard error of the mean
*SNCA*: synuclein alpha gene (encoding α-synuclein)
snRNA-seq: single-nucleus RNA sequencing
STAT: signal transducer and activator of transcription
STING: stimulator of interferon genes
TE: transposable element
TLR: Toll-like receptor
TNF-α: tumour necrosis factor α
TYK2: tyrosine kinase 2
UMAP: uniform manifold approximation and projection
UMI: unique molecular identifier

