## Supplementary Tables, Figures and References for "Astrocyte-Specific dsRNA–ADAR1 Signalling Links Innate Immunity to Complement Activation in Parkinson’s Disease"

**Supplementary Table 1.** Secondary antibodies and counterstaining for the immunofluorescence study

| Secondary antibodies | Company, catalogue number | Concentration |
|---|---|---|
| Donkey anti-Mouse IgG (H+L) Highly Cross-Adsorbed Secondary Antibody, Alexa Fluor™ 488 | Thermo fisher, A21202 | 1:500 |
| Donkey anti-Chicken IgY (H+L) Highly Cross Adsorbed Secondary Antibody, Alexa Fluor™ 568 | Thermo fisher, A78950 | 1:500 |
| Donkey anti-Rabbit IgG (H+L) Highly Cross-Adsorbed Secondary Antibody, Alexa Fluor™ Plus 647 | Thermo fisher, A32795 | 1:500 |
| Alexa Fluor® 790 AffiniPure® Goat Anti-Mouse IgG, Fcγ subclass 2a specific | Jackson Immuno Research Laboratories, 115-655-206 | 1:500 |
| Zenon™ Rabbit IgG Labeling Kit Alexa Fluor 488 | Thermo fisher, Z25302 | Per manufacture |
| Hoechst 33342 | Sigma-Aldrich, B2261 | 1:1000 |

**Supplementary Table 2.** Uniform quality control and preprocessing parameters applied across all datasets for the transcriptomic analyses.

| Parameter | Value | Rationale |
|---|---|---|
| Minimum genes per nucleus | ≥500 | Excludes empty droplets and low-quality nuclei |
| Minimum UMI counts per nucleus | ≥500 | Removes ambient RNA fragments |
| Maximum mitochondrial reads | ≤10% | Filters damaged nuclei; consistent with original studies |
| Upper-bound counts & genes | 99th percentile per sample | Removes likely multiplets and outliers |
| MALAT1 removal | Yes | Nuclear-enriched lncRNA dominating snRNA-seq clustering |
| Ribosomal gene removal (RPL/RPS) | Yes | Uninformative in single-nucleus protocols |
| Doublet detection | scDblFinder per sample | Samples with <100 cells nuclei exempt |

**Supplementary Table 3.** Composition of the nine innate immune sub-panels.

| **Sub-panel** | **Genes (n)** | **Reference** |
|---|---|---|
| ISG core | *ISG15, IFIT1, IFIT2, IFIT3, MX1, MX2, IFITM1, IFITM2, IFITM3, RSAD2, BST2, USP18 (12)* | (1) |
| IFN–STAT | *STAT1, STAT2, IRF1, IRF3, IRF7, IRF9, JAK1, TYK2 (8)* | (2) |
| NF-κB | *NFKB1, NFKB2, RELA, RELB, NFKBIA, IL6, TNF, CCL2, CCL5, CXCL10 (10)* | (3) |
| Complement | *C1QA, C1QB, C1QC, C3, C4A, C4B, SERPING1, CFB, CFH (9)* | (4, 5) |
| OAS–RNase L | *OAS1, OAS2, OAS3, OASL, RNASEL (5)* | (6) |
| Astrocyte reactivity | *GFAP, VIM, SERPINA3, LCN2, CHI3L1, OSMR (6)* | (7) |
| Inflammasome | *NLRP3, NLRP1, AIM2, PYCARD, CASP1, CASP4, IL18, IL1B, GSDMD (9)* | (8, 9) |
| TLR–dsRNA | *TLR3, TICAM1, TICAM2 (3)* | (10) |
| TLR–DAMP | *TLR2, TLR4, MYD88, TIRAP, IRAK1, IRAK4, TRAF6 (7)* | (11, 12) |

### Supplementary Figures

#### Supplementary Figure 1.

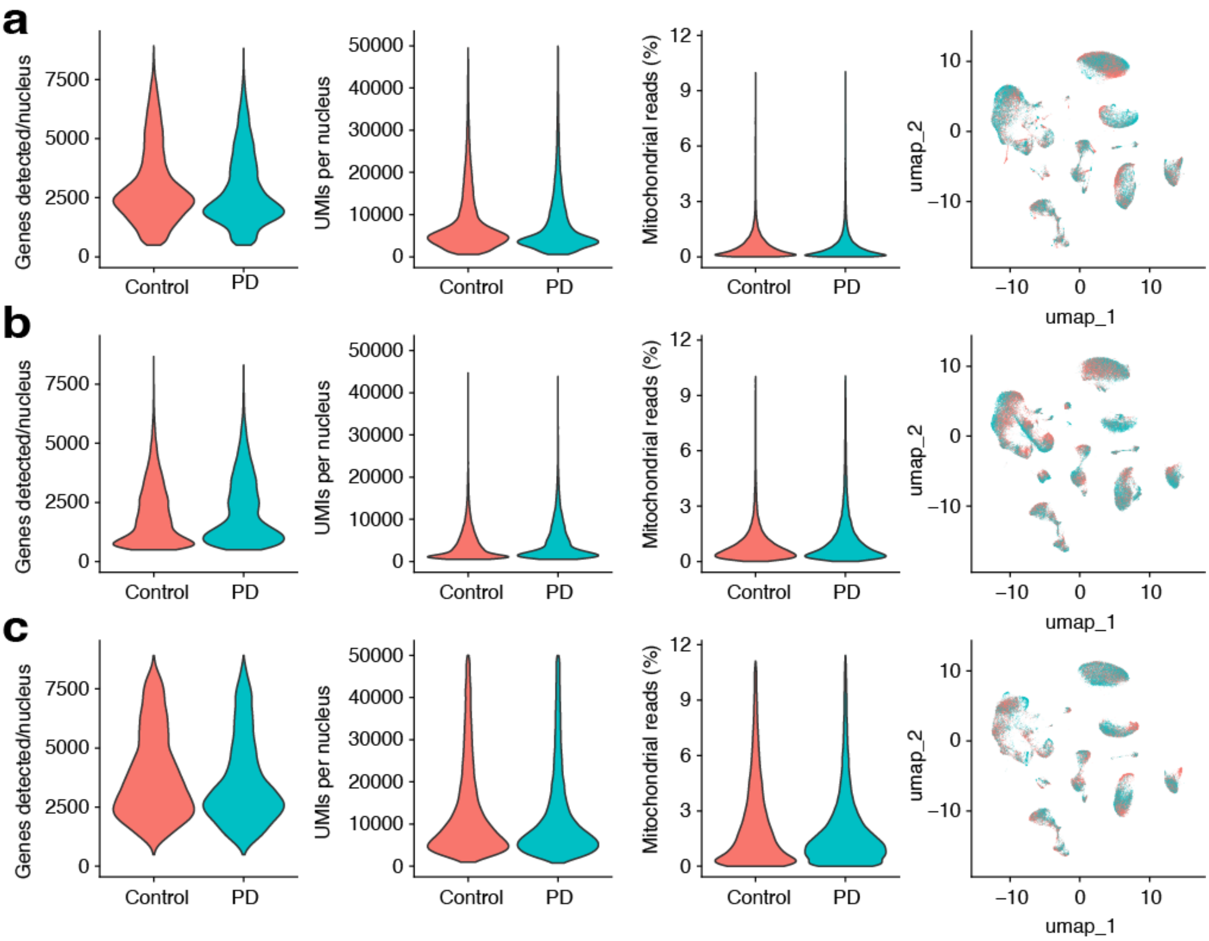


*Per-dataset quality control and dimensionality reduction*. GSE202210 (a), GSE178146 (b), GSE253462 (c) were processed independently using the same prespecified quality-control workflow before cross-dataset integration. Violin plots show the number of detected genes, total unique molecular identifier (UMI) counts and mitochondrial transcript percentage by diagnosis; UMAPs show the pre-integration embedding for each dataset. Each point in a UMAP represents one nucleus.

**Supplementary Figure 2.**

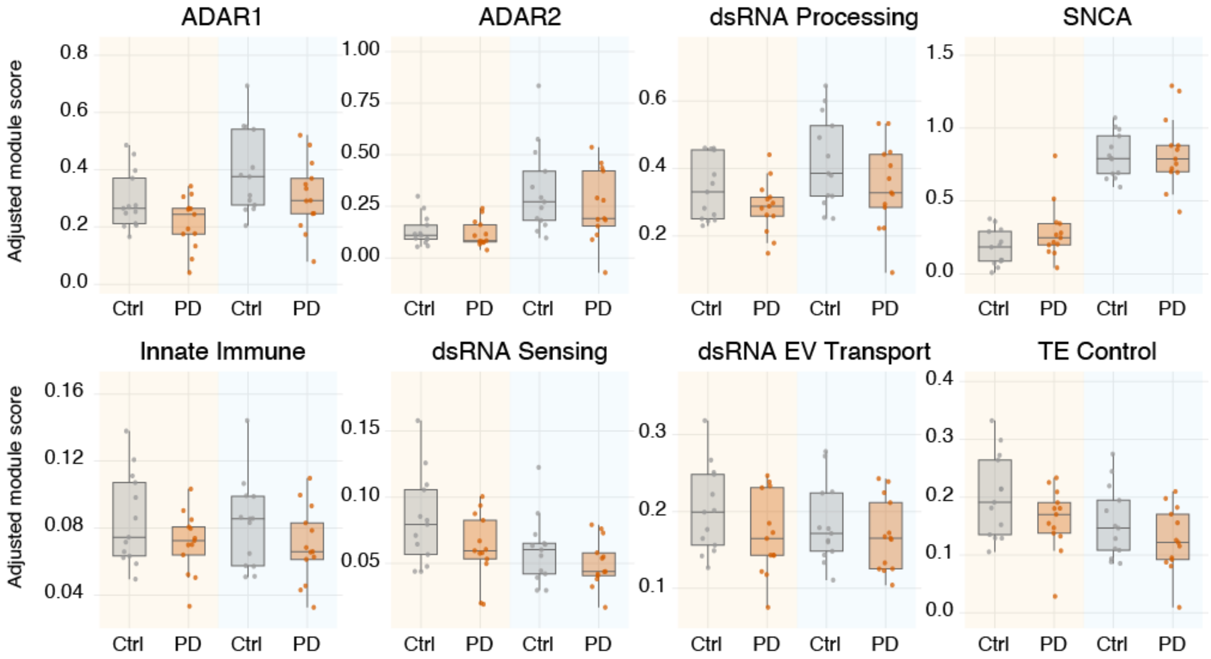


*PD-versus-control comparison of module scores in cortical astrocytes and neurons.* Boxplots show donor-level module scores for ADAR1, ADAR2, dsRNA processing, SNCA, innate immune, dsRNA sensing, dsRNA EV transport and TE control modules. Within each panel, scores are shown separately for astrocytes (cream background) and total neurons (blue background), each split by diagnosis (grey, controls; orange, PD); each point is one donor. BH-adjusted P values are shown for the PD versus control comparison within each cell type. No comparison was significant after correction.

**Supplementary Figure 3.**

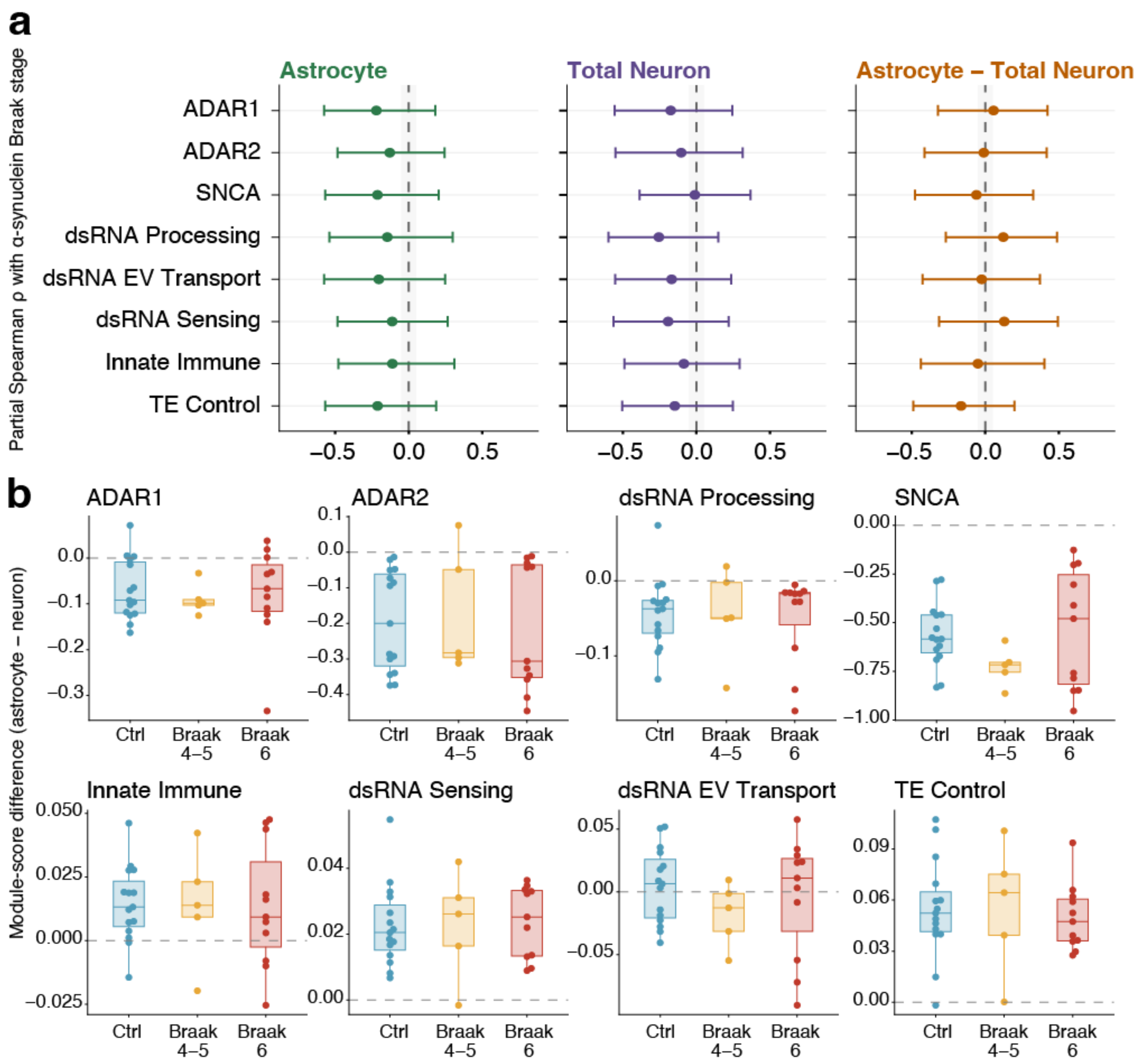


*α-Synuclein Braak stage analysis of cortical neurons and astrocytes*. (a) Partial Spearman correlations between donor-level module scores and α-synuclein Braak stage, shown for astrocytes (green), total neurons (purple) and the astrocyte − total neuron difference (orange). Correlations were adjusted for age and dataset. Points indicate partial Spearman ρ values, and horizontal bars indicate 95% percentile-bootstrap confidence intervals obtained from 10,000 donor-level resamples. The dashed vertical line marks $\rho = 0$, and the shaded band highlights values close to zero ($|\rho| < 0.05$). (b) Within-donor astrocyte–total-neuron module-score differences, calculated as the astrocyte score minus the matched total-neuron score, across control, Braak 4–5 and Braak 6 groups. No Braak-stage association was significant after BH correction across the eight modules within each cell type or divergence analysis.
